# Multivalent Surface Search Dynamics Shape Bacteriophage Adsorption Efficiency: A Stochastic Model of Tail Fiber Optimization

**DOI:** 10.64898/2026.07.03.736286

**Authors:** Anjali Yadav, Kim Sneppen, Namiko Mitarai

**Affiliations:** Niels Bohr Institute, University of Copenhagen, Denmark

## Abstract

Phages must locate and bind to bacterial surface receptors to initiate infection. Their tail fiber configuration critically influences this process. We develop a stochastic model describing surface search as a renewal process, incorporating attachment, detachment, and target-finding steps. Using both numerical simulations and analytical calculations, we quantify how tail fiber number, attachment–detachment rates, and geometric constraints impact the mean and the distribution of time to successful adsorption. Notably, the search efficiency shows a nonmonotonic dependence on tail fiber number, governed by a trade-off between binding stability and diffusion-mediated mobility. This optimum shifts depending on the effective bacterial density, target radius, and fiber reach. Short fiber reach imposes severe geometric constraints, reducing mobility at high tail fiber counts and leading to performance degradation. Our findings suggest that phage adsorption strategies are shaped by a balance between anchoring and exploration, with evolutionary implications for tail fiber design and infection efficiency.

**SIGNIFICANCE:** Bacteriophages typically use tail fibers to bind and explore bacterial surfaces. Using stochastic modeling and numerical simulations, we uncover how attachment-detachment kinetics and tail fiber geometry shape surface search efficiency. Our results reveal a trade-off: more tail fibers stabilize attachment but hinder exploration, especially with short reattachment ranges. These findings offer a mechanistic basis for understanding how phages optimize infection strategies and may explain the diversity of tail architectures observed in nature.

## INTRODUCTION

Bacteriophages, viruses that infect and lyse bacteria, are important players in microbial ecology and evolutionary dynamics (1–3). A critical first step in the phage infection cycle is *adsorption*, the process by which a phage attaches to a bacterial surface and initiates genome delivery. Traditional theoretical models often treat adsorption as an instantaneous, one-step event in which a phage binds to a receptor and immediately injects its DNA (4–6). However, there has been experimental evidence to support the adsorption process involving multiple kinetic processes (7–9). These steps have been explained by models with 3D diffusion of free phages and then 2D diffusion of phages on the bacterial surface after initial binding (7).

Recent single-virus tracking studies have provided direct visualization of the multiple steps of adsorption process (10–14). After initial, reversible contact via one or more tail fibers (15), phages can remain loosely tethered to the bacterial surface and undergo lateral exploration, effectively “walking” across the membrane to search for a suitable receptor. Following reversible attachment and in some cases, surface exploration, phages may proceed to irreversible binding (10–12, 16). This multistep behavior highlights the importance of tail fiber dynamics in determining adsorption efficiency.

Multivalent binding, as a general biophysical principle, has been extensively studied in systems ranging from nanoparticle targeting to receptor-ligand binding (17, 18). In the context of phage biology, this mechanism enables a tunable balance between surface anchoring and mobility—both crucial for efficient infection. Tail fiber number, flexibility, receptor numbers and reattachment range are all hypothesized to modulate this balance (19–22). Bacteriophages display wide variation in tail fiber valency, ranging from single non-contractile tail fiber in some Siphoviridae (e.g., SPP1), through three tail-fibers in Demerecviridae like T5 and four short tail fibers such as real wild type Ur*λ*, to highly multivalent designs with six or more tail fibers or tailspikes as seen in phages T4, T7, P22, and *Listeria* phage A511, reflecting diverse strategies for host recognition and adsorption efficiency (20, 23–29).

Here, we develop a stochastic model to investigate how multivalent binding and tail fiber properties influence phage surface exploration and target finding. We model the adsorption process as a renewal process consisting of cycles of attachment, detachment, and target search. Using numerical simulations and analytical techniques, we examine how key parameters—including the number of tail fibers, their kinetic rates, and spatial reach—affect the mean time to successful adsorption. Our findings reveal a trade-off between surface stability and mobility: while additional tail fibers reduce the risk of detachment, they can also hinder exploratory dynamics, especially when reattachment is spatially constrained. These results provide a mechanistic understanding of how physical constraints shape adsorption and may explain the observed diversity in the architectures of tail fibers between phage species (30, 31).

## MATERIALS AND METHODS

### Stochastic Model Framework

We model the phage as a multivalent stochastic walker that moves via alternating cycles of partial detachment and reattachment Figure 1(a). Each tail fiber can bind to or unbind from the surface at rates *k*_on_ and *k*_off_, respectively, and the overall motion of the phage is governed by the dynamic reconfiguration of the attached tail fibers. We extend the stochastic framework introduced by Kowalewski et al. (30), adapting it to include key biological constraints relevant to phages: a small number of discrete tail fibers (3–10), geometrically constrained reattachment arcs (between nearest attached neighbors) and spatially localized target regions.

**Figure 1:**
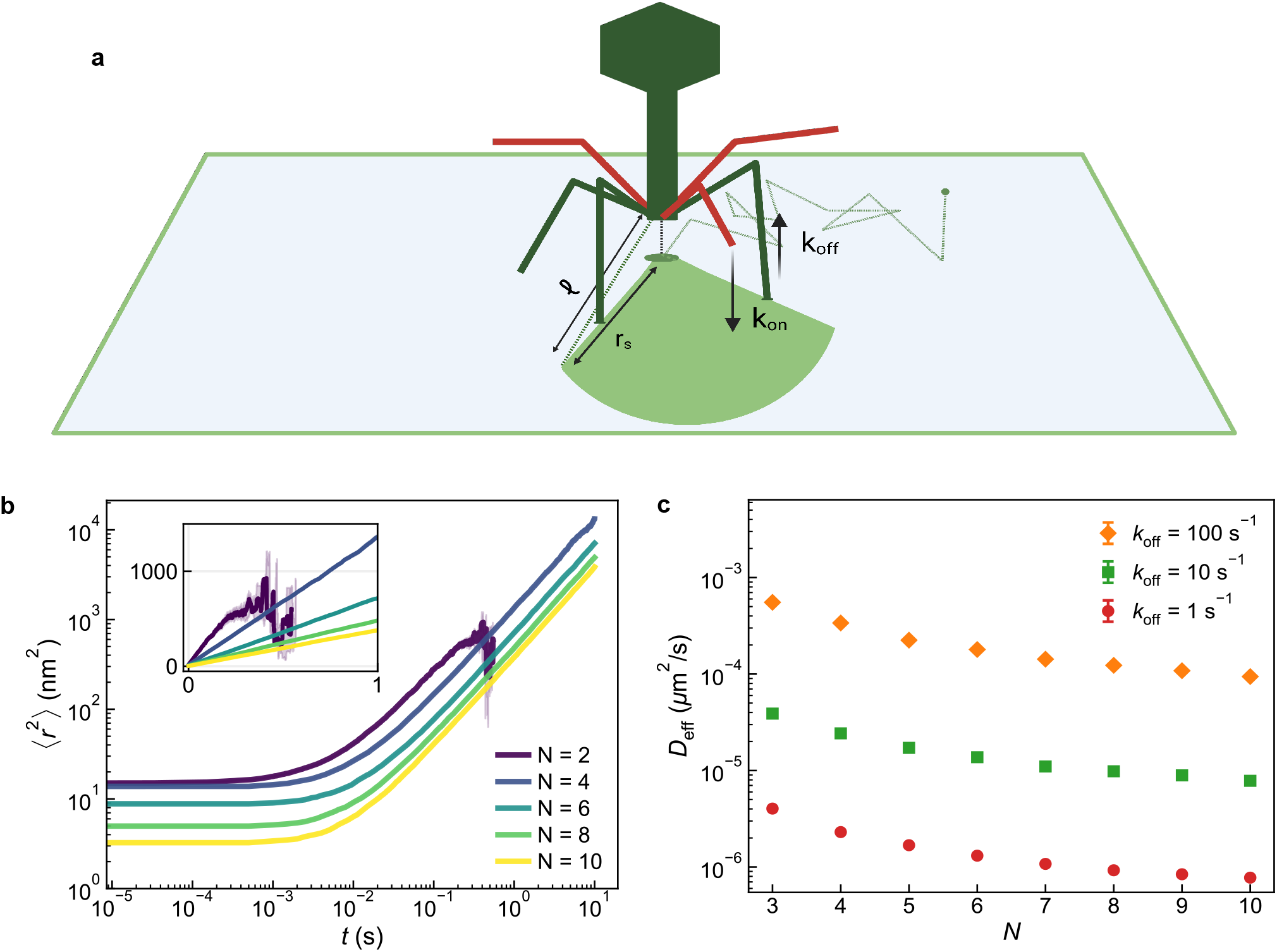
a) Schematic of phage surface exploration in the stochastic model. The phage is represented as a multivalent walker whose tail fibers reversibly attach and detach from a planar surface. Green fibers indicate those currently bound to the surface with detachment rate *k*_off_, whereas red fibers are unbound and can reattach with rate *k*_on_. The light green line traces the diffusion path of the phage center of mass (COM) across the surface, with the green dot directly below marking its instantaneous position, determined by the configuration of all attached fibers. The arrow denotes an unbound (red) fiber about to reattach within the angular sector between two neighboring attached fibers, defining the accessible area for random reattachment. Because receptor binding is not explicitly modeled in diffusion analysis, the surface is treated as homogeneous and receptor-free, allowing diffusion to be governed solely by multivalent geometry and binding kinetics. Created in BioRender. Yadav, A. (2026) https://BioRender.com/d3e6l0g. b) Mean-squared displacement of the phage center-of-mass as a function of time for different tail fiber numbers. (*k*_off_ = 10^2^ s^−1^, *k*_on_ = 10^3^ s^−1^, *r*_*s*_ = 10 nm). MSD is averaged over 1000 trajectories, with shaded regions indicating standard error. The log-log plot emphasizes the early transition before the behavior converges to the ordinary diffusion. The inset is shown in linear scale to visualize the slope. c) Effective diffusion constant *D*_eff_ as a function of tail fiber number *N* for different detachment rates, *k*_off_. *k*_off_ values partially offset the reduction in mobility imposed by multivalency in range *k*_off_ << *k*_on_.

The center-of-mass (COM) of the phage, computed from the positions of the attached tail fibers, defines the locus of motion and detection. After detachment of a tail fiber, reattachment is allowed only within an angular sector between neighboring tail fibers and within a radial distance window of *r*_*s*_. This enforces local exploration and leads to limited lateral movement at high attachment counts, mimicking the realistic loss of freedom in dense multivalent anchoring.

Experimental studies show that bacterial surface receptors can be distributed in helical patterns, clustered, or patch-like depending on the membrane and receptor (32, 33). We chose to instead model receptors as randomly distributed, non-overlapping circular targets of radius *r*_*t*_ on the cell surface. The cell surface is considered a *L*× *L* domain with periodic boundary conditions. Adsorption occurs when the phage COM enters a circular radius receptor zone *r*_*t*_, that may be larger than the actual receptor to take into account the vibrations of the phage within each step of its walk. If all tail fibers detach before this event, the search restarts from a random location, either because the phage diffused back to the same cell surface (which we consider to happen immediately at a fixed probability) (5, 11) or because the phage failed infection and finding a new bacterial surface to infect (which occurs after the diffusion based 3D search). The new search starts from the state where one tail fiber is attached at a random location on the cell surface.

### Renewal Process Formulation

As shown schematically in Figure 2, adsorption can be represented as an iterative cycle of attachment, surface exploration, and detachment until a receptor is encountered. To capture this process quantitatively, we decompose the dynamics into three independent time distributions: the hitting time distribution *P*_*h*_ (*t*), which describes the distribution of the time spent exploring the surface until a receptor is found, starting the search outside the target radius; the detachment time distribution *P*_*d*_ (*t*), which describes the distribution of times until complete loss of attachment when all tail fibers unbind; and the search time distribution *P*_*s*_ (*t*), which reflects the distribution of waiting time before a new host is encountered.

**Figure 2:**
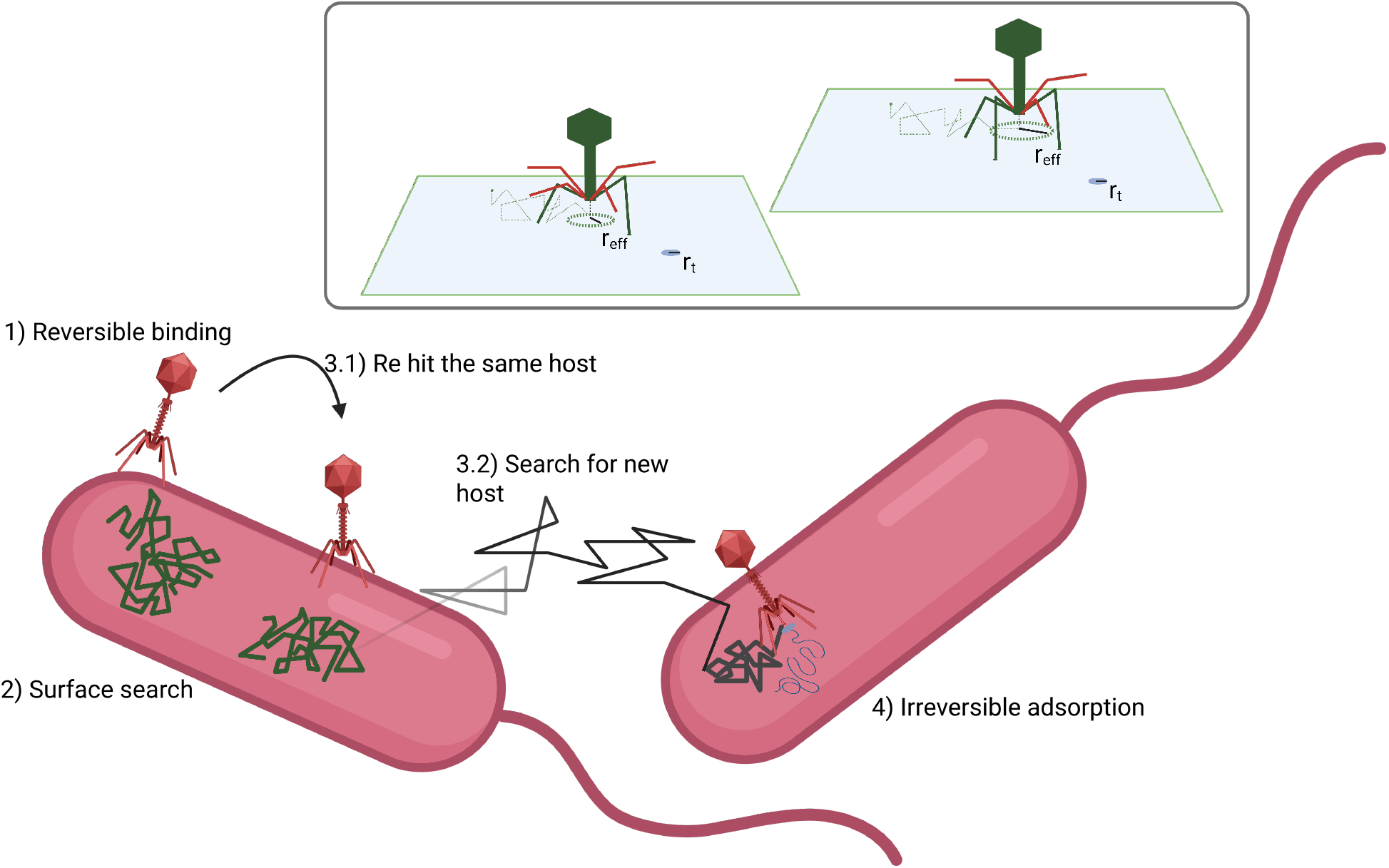
Schematic of the stochastic adsorption process. 1) A phage initially binds reversibly to the bacterial surface via tail fiber. 2) While attached, the phage undergoes lateral surface exploration, modeled as effective diffusion of the center-of-mass with diffusion constant *D*_eff_. 3) Complete detachment occurs when all fibers unbind, after which the phage either reattaches to the same surface with probability *p*_*r*_, or enters bulk diffusion to search for a new host with encounter rate η*B*. 4) Adsorption occurs when the phage encounters a receptor region, leading to irreversible binding. Inset (boxed panel): The effective target size *r*_eff_, due to thermal fluctations of the phage’s center of mass is shown. With fixed set of attached tail fibers, thermal fluctuations allow the phage to explore a finite region around its mean position, effectively enlarging the receptor size. The quantitative impact of *r*_eff_ is discussed in results section. Created in BioRender. Yadav, A. (2026) https://BioRender.com/nod6b1m.

The search time distribution (*P*_*s*_ (*t*)) depends strongly on the bacterial density in the environment: in crowded conditions such as in the gut or well-grown laboratory cultures, reattachment is rapid, whereas in sparse environments, it is slow. In the model, we assume mass action in the collision of a bacterium and a phage, hence *P*_*s*_ (*t*) follows an exponential distribution with mean 1 / (η*B*), where η is the diffusion limited encounter rate per bacterium in a well mixed system and *B* is the bacterial density. To analyze the coupled processes of the total adsorption time distribution *P*_ads_ (*t*), we cast the adsorption problem as a stochastic resetting problem (34). At *t* = 0, a phage starts on the cell surface, with one of the tail fibers attached. Adsorption can occur during surface exploration or after one or more detachments and reattachments. In addition to detachment, two geometric effects are also considered:

- First, after detachment, the phage can reattach to the same cell with the probability of rehitting *p*_*r*_, reflecting the repeated return to the surface of the same cell. Such multiple surface encounters arise naturally from diffusion near partially absorbing boundaries (5).
- Second, upon landing on a cell surface the phage may immediately adsorb with probability 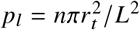 if we assume *n* non overlapping receptors.

*P*_ads_ (*t*) represents the distribution of the total time required for a phage to irreversibly bind a receptor after repeated cycles of attachment, exploration, detachment and reattachment, expressed in the renewal equation (35–38)

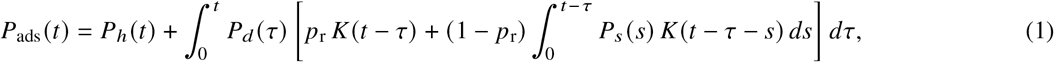

where

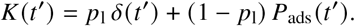

Here, the δ-term represents immediate adsorption upon landing, while the second term accounts for restarting the adsorption process when the landing position lies outside the target region. For exponential waiting times, Equation (1) is equivalent to a set of linear rate equations (Supplemental Section S1).

### Parameter Selection and Biological Context

All model parameters are expressed in dimensionless units. This allows systematic exploration of parameter space while allowing translation to real values as summarized in Table 1. The fundamental scaling units and parameter values were chosen based on the literature on bacteriophage-host interactions as described in the following paragraphs.

**Table 1:**
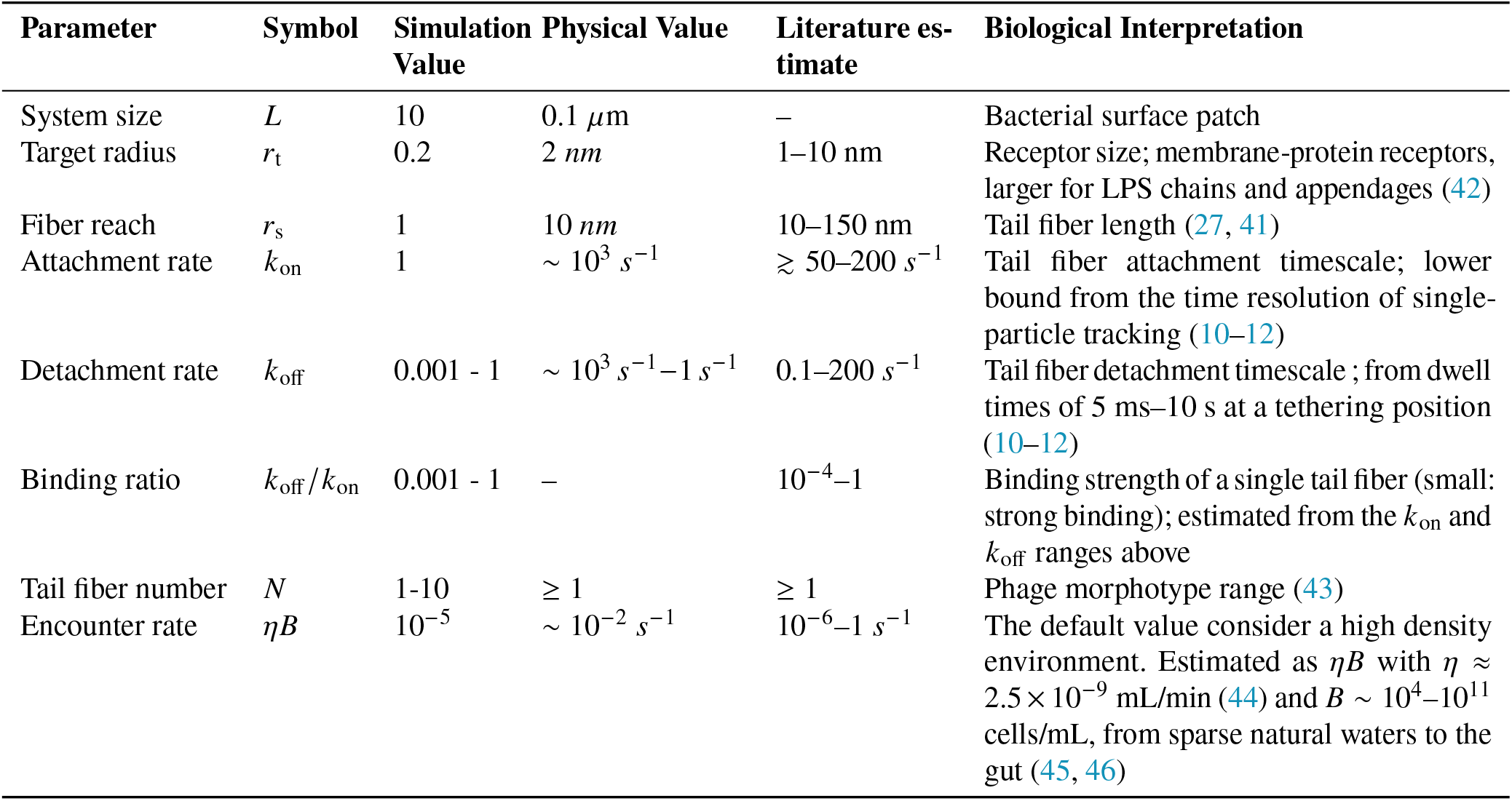
Summary of model parameters with physical interpretations and literature basis. Physical values are provided for order of magnitude mapping only; all simulations are performed in dimensionless units.

| Parameter | Symbol | Simulation Value | Physical Value | Literature estimate | Biological Interpretation |
| --- | --- | --- | --- | --- | --- |
| System size | $L$ | 10 | $0.1 \mu\text{m}$ | – | Bacterial surface patch |
| Target radius | $r_t$ | 0.2 | $2 \text{ nm}$ | 1–10 nm | Receptor size; membrane-protein receptors, larger for LPS chains and appendages (42) |
| Fiber reach | $r_s$ | 1 | $10 \text{ nm}$ | 10–150 nm | Tail fiber length (27, 41) |
| Attachment rate | $k_{\text{on}}$ | 1 | $\sim 10^3 \text{ s}^{-1}$ | $\gtrsim 50\text{--}200 \text{ s}^{-1}$ | Tail fiber attachment timescale; lower bound from the time resolution of single-particle tracking (10–12) |
| Detachment rate | $k_{\text{off}}$ | 0.001 – 1 | $\sim 10^3 \text{ s}^{-1} \text{--} 1 \text{ s}^{-1}$ | $0.1\text{--}200 \text{ s}^{-1}$ | Tail fiber detachment timescale; from dwell times of 5 ms–10 s at a tethering position (10–12) |
| Binding ratio | $k_{\text{off}}/k_{\text{on}}$ | 0.001 – 1 | – | $10^{-4}\text{--}1$ | Binding strength of a single tail fiber (small: strong binding); estimated from the $k_{\text{on}}$ and $k_{\text{off}}$ ranges above |
| Tail fiber number | $N$ | 1–10 | $\geq 1$ | $\geq 1$ | Phage morphotype range (43) |
| Encounter rate | $\eta B$ | $10^{-5}$ | $\sim 10^{-2} \text{ s}^{-1}$ | $10^{-6}\text{--}1 \text{ s}^{-1}$ | The default value consider a high density environment. Estimated as $\eta B$ with $\eta \approx 2.5 \times 10^{-9} \text{ mL/min}$ (44) and $B \sim 10^4\text{--}10^{11} \text{ cells/mL}$, from sparse natural waters to the gut (45, 46) |

The system is assigned a linear size *L* = 10 (dimensionless units) corresponds to a local patch of bacterial cell surface. In physical units, this corresponds to a surface of approximately 0.1 µm ×0.1 µm. The length scale is sufficient to capture the localized surface exploration dynamics. Experiments show that bacteriophages explore confined regions of the host’s surface, rather than uniformly sampling the entire cell surface (10–13, 39).

In physical units, the target size *r*_t_ = 0.2 corresponds to approximately 2 nm, consistent with the nanometer-scale dimension of bacteriophage receptors on bacterial surfaces, typically of order 1-10 nm (40). The tail fiber reach, *r*_s_ = 1.0, corresponds to 10 nm in physical units. We later vary this parameter to reflect the diversity of tail fiber architectures in phage families, from the extended fibers of T4 (145 nm) to the shorter fibers of T7 (10–20 nm) (27, 41).

It should also be noted that thermal fluctuation of the phage affects their effective search radius of the target when the attachment of tail fibers do not change (Figure 2 inset). Such fluctuation can make the target size of the receptor effective larger. A simple estimate based on an entropic spring model of tail fibers shows that the effective target search radius can easily be on the order of 10nm (Supplemental Figure S1). Although we initially set the default target radius to be the receptor protein radius (*r*_*t*_ = 0.2), we later discuss how the results are modified when the effective target radius is larger.

Direct measurements of single tail fiber binding and unbinding rates are limited. However, single-cell tracking experiments show that the bacteriophage undergoes repeated reversible attachment events on sub-second time scales during surface exploration (10–12). High-speed fluorescence microscopy of bacteriophage T4 reveals discrete step-like motion near/on the cell; Analysis of these step durations shows that shifts between successive positions occur at rates equal to or faster than the temporal resolution of the experiment (5-20 ms), indicating that underlying attachment-detachment events must occur on submillisecond timescales (11). This constrains the microscopic binding unbinding rates to be larger than 100/ sec - 1000/ sec. In this model, time units are dimensionless by setting *k*_on_ = 1, which thereby defines our timescale. Consequently, all other rates are relative to this attachment timescale. The detachment rates (*k*_off_) varied between 0.001 and 1.0 (in units of *k*_on_), corresponding to the residence times (τ = 1/*k*_off_) between 1 and 1000 in units of the attachment time.

The bacterial encounter rate η*B* was set to 10^−5^ in dimensionless units, unless otherwise noted. This parameter represents rare effective encounters between free phages and host cells and is determined by the bacteria density in bulk. Higher values of η*B* correspond to high-abundance environments such as laboratory cultures or the gut, while lower values represent low-abundance environments such as the open ocean (47). We explore different values of η*B* to consider the effect of bacterial concentration in different environments.

The above parameters are used in all simulations, unless stated otherwise. Simulations were performed using the Gillespie algorithm with the following specifications: 5000 independent trajectories per parameter set, maximum simulation time of 10,000 dimensionless units. The number of tail fibers was varied from 2 to 10, covering the typical diversity found among tailed phages (43). Unless otherwise stated, we consider a single receptor target on the surface. When multiple targets are considered, they are assumed to be randomly distributed without overlap.

## RESULTS AND DISCUSSION

### Increased tail number constrain surface mobility

To quantify how tail fiber number affects phage surface exploration, we first measured the mean-squared displacement (MSD) of the phage center-of-mass during surface-bound diffusion. This is done without considering the adsorption sites. Simulations uses baseline dimensionless parameters *k*_off_ = 0.1, *k*_on_ = 1.0, and *r*_s_ = 1. The initial condition is set by one tail fibers attached to the cell surface.

The MSD curves Figure 1(b) exhibit two distinct temporal regimes: an initial sub-diffusive plateau reflecting configurational equilibration, followed by a crossover to normal diffusion with slope 4*D*_eff_. The crossover time scales with *N*, indicating that geometric constraints slow the approach to steady-state diffusion. However, this transient is brief compared to the diffusive phase, allowing us to extract long-time diffusion coefficients.

The effective diffusion constant *D*_eff_ decreases systematically with the number of tail fibers, both for low and high detachment rates, as seen in Figure 1(c). The scaling follows approximately *D*_eff_ ∝*N*^−α^ with α ≈1. The exponent 1 can be understood from simple geometric considerations. The number of tail fibers divides the circumference of the phage into *N* sectors of the central angle Δθ∼ 2π/ *N*. As a result, the tail fiber reattachment event of a length *r*_*s*_ displaces the center of mass of the phage by Δ*r* = *r*_*s*_Δθ. Each step happens if any of the *N* tail fibers attach or detach, and would occur at the mean rate, *N k*_off_ *k*_on_/(*k*_off_ + *k*_on_). In the limit, *k*_off_ << *k*_on_, i.e, when phage is stable on the surface to explore, effective diffusion constant scales as 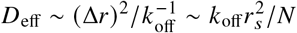. The steeper dependence observed for *k*_off_ ∼ *k*_on_ reflects deviations from this limiting scaling as the full expression for *D*_eff_ becomes relevant Figure 1(c).

Consistent with this interpretation, increasing *k*_off_ shifts the entire *D*_eff_ (*N*) curve upward (Figure 1(c)). A higher detachment rate increases the frequency of attachment–detachment events while also reducing the average number of simultaneously attached tail fibers. As a result, the geometric constraints imposed by multivalent binding are partially relaxed, allowing larger displacements of the phage center of mass and therefore faster surface diffusion. This effect becomes more pronounced as *k*_off_ approaches *k*_on_, where multiple tail fibers are frequently detached.

Similarly, a larger tail fiber reach *r*_s_ enhances mobility, following an approximate scaling 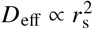, as shown in Figure 5(c).

Longer tail fibers increase the distance over which reattachment can occur, thereby increasing the displacement associated with each reattachment event. Consequently, the phage explores a larger area of the surface during each attachment–detachment cycle. This geometric advantage partially compensates for the reduced mobility caused by additional tail fibers, suggesting that longer tail fibers can offset the effects of reduced diffusion due to increased multivalency and maintain efficient surface exploration.

### Hit-time statistics follow exponential kinetics with diffusion-controlled rates

Once the phage is attached to the bacterial surface, it either locates directly a receptor within the range *r*_*t*_ of the first attachment or searches the host surface through diffusion. Here, we consider the search where it diffuses until it finds the target receptor or detaches from the surface. For analytical evaluation of this search time, we consider the surface search as diffusion of the COM at a diffusion coefficient *D*_eff_ on a periodic *L*× *L* domain with a small circular absorbing receptor of radius *r*_*t*_ ≪*L*. For now we ignore the detachment event, which will be discussed next. After initial transient, on a finite periodic domain, the target encounter rate becomes constant, and the hitting time distribution is well approximated by an exponential 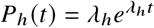 with rate *λ*_*h*_ = 1/ ⟨*T*_*h*_⟩ (36).

For a diffusing particle with a diffusion constant *D*_eff_ in a periodic domain, the mean first-passage time to find one target of radius *r*_*t*_ on the 2-D surface is (48)

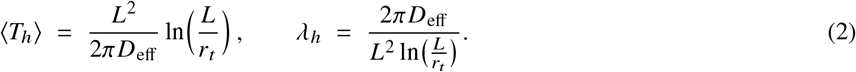

The derivation, adapted to our notation and geometry, can be found in Supplemental Section S5 (36). Simulations confirm that the hit time distribution *P*_*h*_ (*t*) is well described by 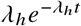 for all parameter sets with sufficient statistics, and the fitted *λ*_*h*_ agrees with Eq. (2) when *D*_eff_ is taken from MSD (Figure 3(a)).

**Figure 3:**
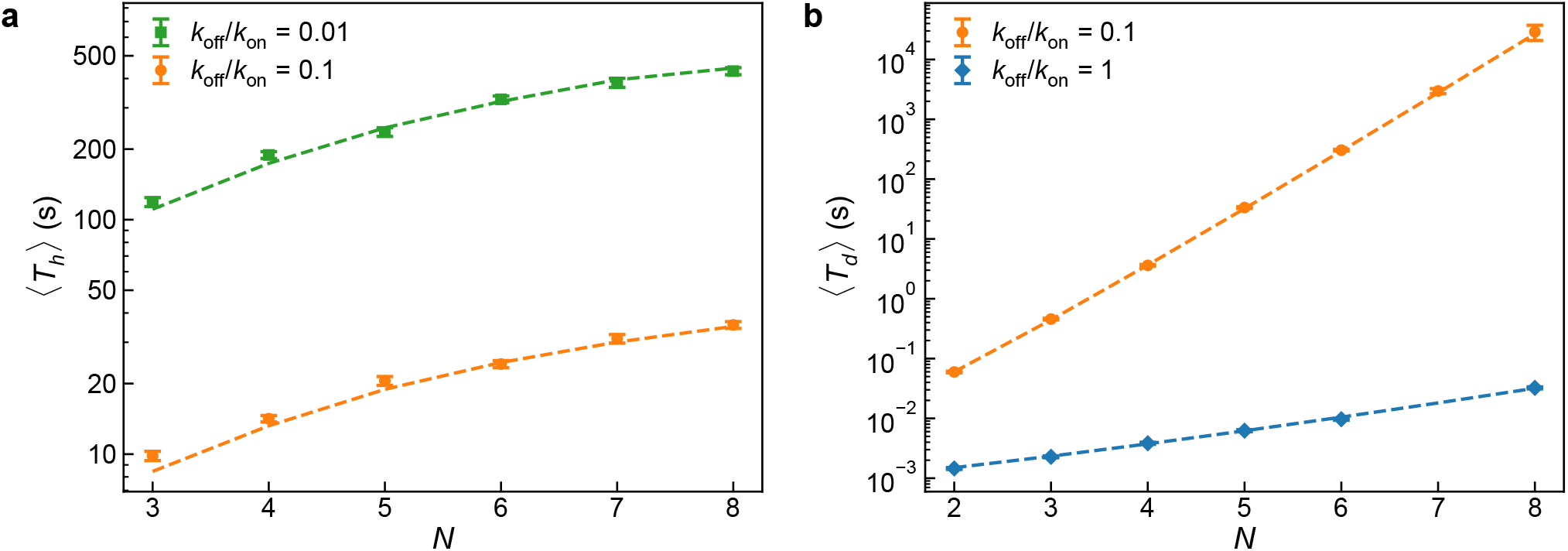
a) Comparison between theoretical and simulated hitting times; the dashed line corresponds to the theoretical prediction, ⟨*T*_*h*_⟩ b) Comparison between theoretical and simulated detachment times; the dashed line corresponds to the theoretical prediction, ⟨*T*_*d*_ ⟩, and the scatter points with error bars show the mean detachment time for 1000 simulations and error.

This analysis can be extended to multiple targets case (Supplemental Section S5). For *n* well-separated receptors 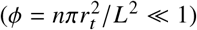, the dilute limit mean hit time generalizes to

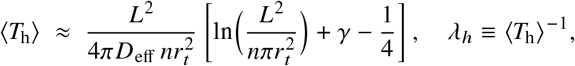

with the Euler’s constant γ = 0.5772; *P*_*h*_ (*t*) remains approximately exponential with rate *λ*_*h*_.

Simulation-derived hit-time distributions closely follow this exponential form throughout the full dynamic range. The fitted rates *λ*_*h*_ agree quantitatively with theoretical predictions based on measured diffusion coefficients as shown in Figure 3(a), confirming that surface search is indeed diffusion-limited under our geometric conditions.

### Detachment kinetics obey birth-death first-passage statistics

Although surface diffusion facilitates the search for a receptor, successful adsorption also requires that the phage stay attached to the bacterial surface long enough to complete this search. The complete loss of surface contact occurs when all *N* tail fibers detach before any can reattach. We model this process as a continuous-time birth–death process in which the number of fibers of the bound tail *n* (*t*) evolves with detachment transitions *n* →*n*− 1 at rate *λ*_*n*_ = *k*_off_*n* and reattachment transitions *n* →*n*+ 1 at rate µ_*n*_ = *k*_on_ (*N* −*n*). This Markov process yields a closed system of equations for the state probabilities *P*_*n*_ (*t*) (Supplemental Section S4). The complete detachment time corresponds to a first-passage event from any state *n* ≥1 to the absorbing boundary at *n* = 0 and the reflecting boundary condition at *n* = *N*.

The effective detachment rate entering the renewal equation is given by the inverse mean first-passage time starting from a single attached tail fiber,

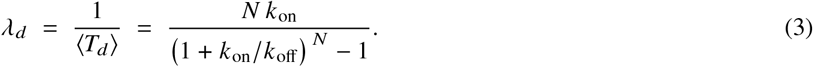

This reduces to ⟨*T*_*d*_⟩∝ (*k*_on_ /*k*_off_)^*N* −1^ /*k*_off_ in the strong-binding regime, where the mean detachment time grows exponentially with tail fiber number, reflecting the stabilizing effect of multivalent binding. The simulation of mean detachment-time agrees quantitatively with the theoretical predictions of birth–death across all combinations of parameters tested (Figure 3(b)).

### Combined renewal model reproduces total adsorption kinetics

Using the validated component distributions—exponential *P*_*h*_ (*t*) with rates determined from simulated *D*_eff_ values from Figure 1, birth-death *P*_*d*_ (*t*) from the attachment model rates using Equation (3), and exponential host encounter *P*_*s*_ (*t*) with mean 1/ (η*B*) —the renewal equation Equation (1) that quantitatively reproduces the simulated total adsorption-time distributions (Figure 4(a)). The biphasic distribution reflects two characteristic classes of adsorption trajectories. The initial fast component corresponds to phages that successfully locate the receptor during their first surface encounter, before complete detachment occurs. Consequently, the majority of adsorption events take place on this short timescale. The slower component arises from trajectories in which the phage detaches before locating the receptor and must undergo one or more additional attachment–detachment cycles, including bulk search for the same or a new host, before successful adsorption. The separation of these timescales therefore reflects the competition between successful surface exploration and stochastic resetting through complete detachment.

**Figure 4:**
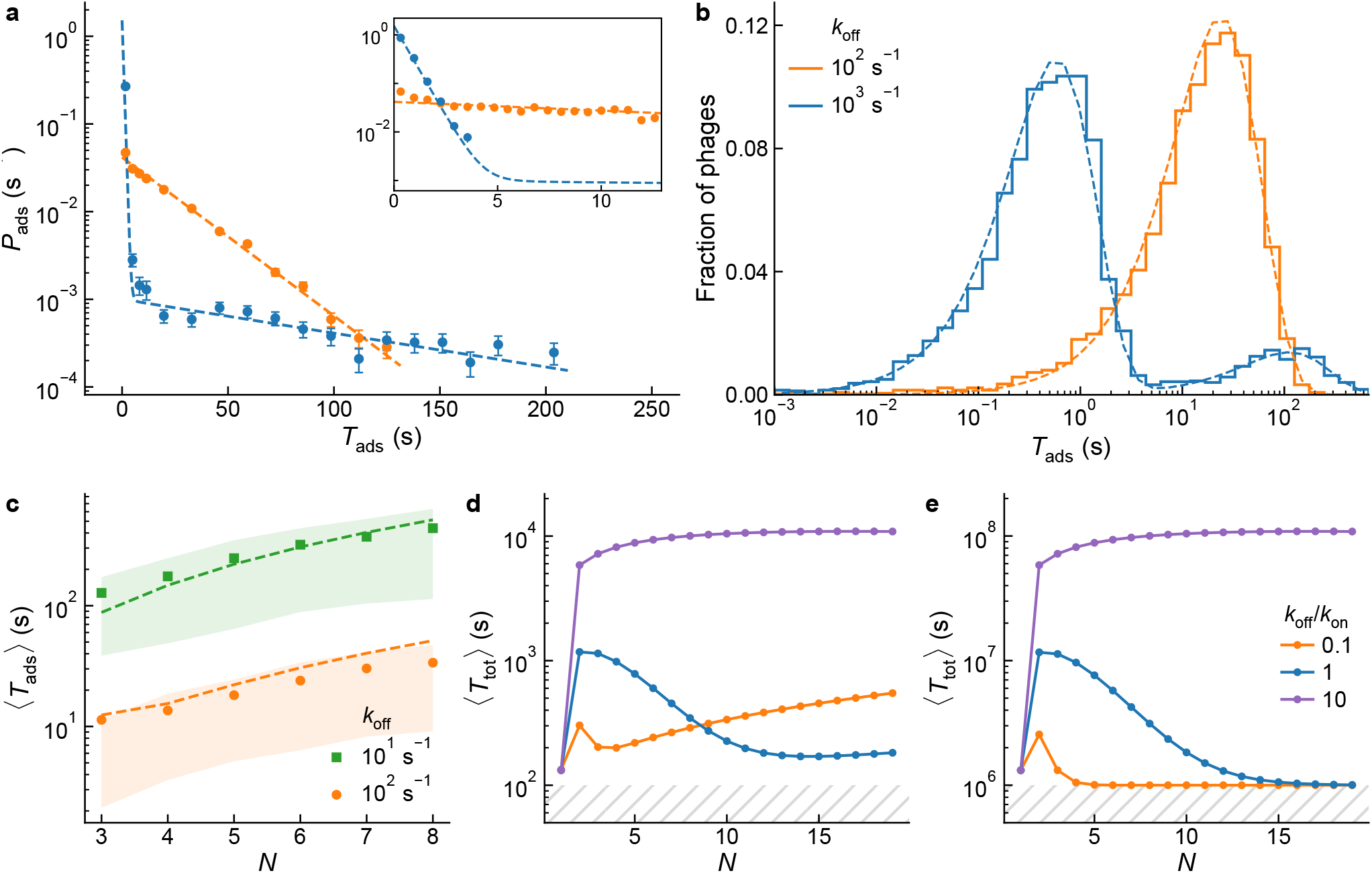
a) Adsorption time distribution, *P*_ads_ (Start from one tail fiber attached to a cell surface, *N* = 6). Dashed lines show theoretical distribution from renewal model Equation (1), using the hitting rate, *λ*_*h*_ and detachment rate, *λ*_*d*_, estimated from the simulated surface cycles. Points show simulation result for different detachment rate, *k*_off_. Inset: the early decay (first∼ 13 s) on the same axes. b) Same data as in (a), shown as the fraction of phages adsorbing per logarithmic time bin. c) Points show the mean adsorption time ⟨*T*_ads_⟩ obtained from stochastic simulations as a function of tail fiber number *N*, for two detachment rates *k*_off_ = 10 s^−1^ and 10^2^ s^−1^. Shaded regions indicate the interquartile range (25–75%) of the adsorption-time distribution. Solid dashed lines show the theoretical prediction from the renewal framework Equation (4). Simulated adsorption times include contributions from repeated attachment–detachment cycles and stochastic bulk search after detachment, with reattachment to the same surface occurring with probability *p*_*r*_ = *a*/(*a* +*r*_*t*_) after each detachment. d) Mean total adsorption time ⟨*T*_tot_⟩ = ⟨*T*_ads_⟩+ 1 / (η*B*) as a function of *N* for high density environment (η*B* = 10^−2^ s^−1^). e) Mean total adsorption time ⟨*T*_tot_⟩ as function of *N* for low density environment (η*B* = 10^−6^ s^−1^) for different ratios *k*_off_/ *k*_on_. Parameters: *k*_on_ = 10^3^ s^−1^, *r*_*s*_ = 10 nm, *r*_*t*_ = 2 nm, η*B* = 10^−2^ s^−1^, and a single target; five targets in (d, e). Times are in seconds. Panels (d, e) show the total adsorption time ⟨*T*_tot_⟩, measured from the instant a phage begins searching in the bulk until receptor binding. Thus, unlike panels (a–c), the total adsorption time includes both the bulk search time, which depends on the bacterial density, and the subsequent surface exploration time, given by ⟨*T*_ads_⟩.

The excellent agreement in Figure 4(a) confirms that the overall adsorption kinetics emerge from the interplay between diffusion-limited surface search, multivalency-controlled detachment resistance, and environmental encounter statistics. Importantly, this framework enables prediction of total adsorption times from independently measurable parameters such as single-particle dwell times and on-cell motion(11, 12, 14).

### Environmental and geometric parameters determine optimal tail fiber number

The complete adsorption process depends critically on the number of tail fibers, which determines both surface mobility and binding stability. To understand how phages optimize this architectural parameter, we analyze the mean adsorption time as a function of the number of tail fibers *N* under varying environmental and kinetic conditions.

For *N* ≥2, from the renewal framework Equation (1), combined with the measured scaling *D*_eff_ (*N*) ∝ *N*^−α^ given in Figure 1(c) and the detachment rate from birth-death process, and the probabilities of same-cell re-encounter and direct landing on the target, the mean adsorption time (Supplemental Section S6) takes the approximate form

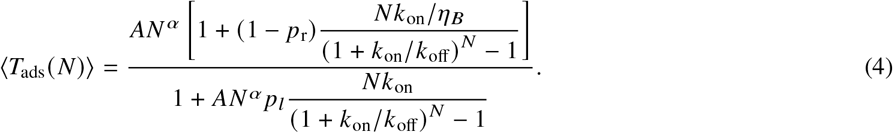

Where *p*_*r*_ is the probability of phage reattachment to a bacteria after leaving, and *p*_*l*_ the probability of direct landing and binding to a receptor at the time of reattachment. α≈ 1 is the measured diffusion penalty exponent (Figure 1(c)), and the prefactor *A* collects geometry-dependent constants and baseline kinetic factors (see Supplemental Section S5).

The resulting mean adsorption time ⟨*T*_ads_ (*N*) ⟩ (Equation (4)) is compared with simulations in Figure 4(c). Theoretical predictions show excellent agreement with the simulations explored in *N*, validating the renewal-based formulation. The case *N* = 1, would be treated separately, as the surface walk would no longer be applicable. In this limit, detachment occurs with rate *k*_off_, and the mean adsorption time is instead

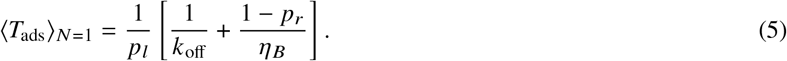

Phage *λ* provides a test case for this limit. The common laboratory strain carries a single central tail fiber (Table 3), and in laboratory culture it lies in the high-density regime where Figure 5(a) predicts a small optimal valency. The quantities entering Equation (5) have been measured for this system. The encounter rate constant is η≈ 2 ×10^−11^ cm^3^ s^−1^, and the host carries ≈280 LamB receptors, a surface density of ≈30 µm^−2^ (9), within a factor of a few of the density of one receptor per *L*× *L* patch used as our default, and the mean on-cell dwell time of≈ 1 s (10) places an upper bound *k*_off_ ≲ 1 *s*^−1^ on the detachment rate of its single fiber. These values are consistent with the parameters used here.

**Figure 5:**
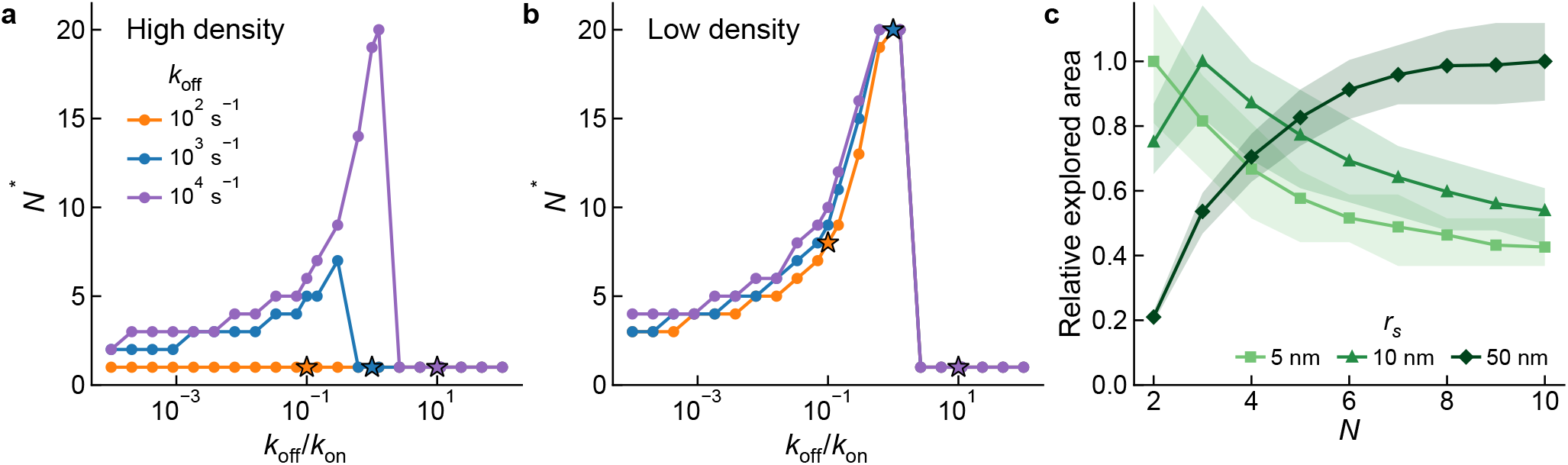
Optimization of tail fiber number for total adsorption time. a) for high density η*B* = 10^−2^ s^−1^, b) for low density η*B* = 10^−6^ s^−1^, as a function of *k*_off_ /*k*_on_ for different *k*_off_ values. The optimal tail fiber number is obtained by minimizing ⟨*T* (*N*) ⟩ over *N*∈ [1, 20], with *N* treated as a discrete variable. Starred points (⋆) correspond to the parameter pairs *k*_off_ /*k*_on_ = 0.1, 1.0, and 10.0 whose full ⟨*T*_tot_ (*N*) ⟩ curves are shown in Figure 4(d, e). c) Relative explored area as a function of *N* for *r*_*s*_ = 5, 10 and 50 nm: number of distinct surface sites (2× 2 nm^2^ lattice) visited by the phage center of mass within 2 s, divided by the maximum of each curve. Simulations at *k*_off_/ *k*_on_ = 0.1, using only trajectories that remain attached during the whole time window; lines show the mean and shaded regions the 25–75% range over trajectories.

The assumption of no surface walk at *N* = 1 is nevertheless an idealization. Phage *λ* shows two-dimensional diffusion on the host surface before irreversible binding (10). One possible origin of this motion within our framework is repeated detachment and reattachment of phage to the same cell, which produces a correlated sequence of contact points that would appear as continuous diffusion at the time resolution of the experiment, corresponding to the limit *p*_*r*_ →1 in Equation (1). The displacement per step in this case would be set by the distance traveled between unbinding and landing rather than by the arc *r*_*s*_Δθ of Figure 1(a). The time resolution is not sufficient to establish the mechanism, and the motion could equally arise without full detachment, for example if the phage moves around a single attached fiber or through the weak reversible receptor interaction proposed in (10). In either case the mobility of surface attached phage with *N* = 1 is finite rather than zero, hence Equation (5) somewhat overestimates ⟨*T*_ads_⟩. This would further favor low valency in dense environments.

To explore the dependence of adsorption efficiency on environmental conditions Figure 4(d, e) show the mean total adsorption time ⟨*T*_tot_⟩ when the phage start from a free, unattached state. This will add the time for the phage to encounter a cell surface first, hence ⟨*T*_tot_⟩ = ⟨*T*_ads_ (*N*) ⟩+1 / (η*B*). This is calculated from Equation (4) and Equation (5) over a wider range of *N* in two representative regimes of host density.

In high-density environments Figure 4(d), detachment is less costly because a detached phage rapidly encounters another host. The dominant competition here is between two strategies, detaching and landing successfully on the receptor on a new surface or remaining attached and searching the surface for the receptor by diffusion. Since new encounters are frequent, *N* = 1 strategy can be very efficient. Introducing additional tail fibers shifts the dynamics towards surface search, but the attachment lifetime is too short to reliably find the receptor before detachment, and the phage pays both the search cost and detachment cost, hence increasing the adsorption time. At larger *N*, the attachment becomes stable enough to complete the surface search and find receptor decreasing *T*_ads_, but since the penalty for fully detaching and landing successfully on the receptor is small, the benefit of stability on the surface is much weaker than in the case of low density, and the surface search strategy rarely outperforms the repeated landing strategy.

In low-density environments Figure 4(e), host encounter after detachment is slow. Therefore detachment is expensive. For *N* = 1, adsorption occurs mainly through repeated direct landing attempts on the receptor, which is inefficient when *p*_*l*_ << 1, i.e, receptor density is low. Adding tail fibers first slows surface diffusion, so ⟨*T*_ads_⟩ increases. However, at larger *N*, the gain in attachment lifetime increases and the phage remains on the surface long enough to find the receptor before detaching. After that, increasing *N* limits the surface diffusion. However, the attachment lifetime is sufficient enough to explore the surface. This produces the decrease and plateau in Figure 4(e). The position of this plateau depends on the *k*_off_/ *k*_on_ ratio; weaker binding requires more fibers before stable surface search becomes effective, whereas stronger binding reaches this regime at smaller *N*. Figure 5 summarizes these observations in terms of the optimal tail fiber number *N*^∗^ that minimizes the mean total adsorption time ⟨*T*_tot_⟩ for a given host density, The value of *N*^∗^ was obtained by discrete numerical minimization over integer values of *N* up to 20, and plotted as a function of the kinetic ratio *k*_off_/ *k*_on_ for two regimes of bacterial density representing dense laboratory cultures (η*B* = 10^−5^); Figure 5(a) and sparse natural environments (η*B* = 10^−9^); Figure 5(b). *k*_off_ /*k*_on_ > 1 corresponds to weak effective binding and frequent loss of contact with the surface, whereas *k*_off_ /*k*_on_ < 1 corresponds to stable attachment. The absolute rates (the values of *k*_off_), shown in different colors, determine the timescale on which individual tail fiber attachment-detachment is carried out, and hence affect the effective diffusion.

The effective diffusion is especially important in high density environments, where surface exploration directly competes with the time required to detach and encounter another host. If the attachment-detachment dynamics is too slow, the repeated landing strategy then remains competitive over a wide range of *k*_off_/ *k*_on_, keeping *N*^∗^∼ 1. If the attachment–detachment dynamics is faster, the phage can diffuse and explore the surface before a full detachment–attachment to a new host cycle starts, and the multivalent surface search becomes competitive at moderate *N*. This is visible in Figure 5(a), where separate curves for different absolute *k*_off_ values differ significantly at intermediate *k*_off_/ *k*_on_.

In the low-density regime Figure 5(b), where detaching and then needing to find another host becomes the significant cost, the dependence on absolute rates becomes weaker. Across all tested *k*_off_, we observe a non-monotonic dependence of *N*^∗^ on *k*_off_/ *k*_on_: at small *k*_off_ /*k*_on_ (stable attachment), *N*^∗^ takes moderate values and increases gradually as *k*_off_ /*k*_on_ grows, since more tail fibers are needed to compensate for the increasing detachment cost. *N*^∗^ reaches a maximum near *k*_off_ /*k*_on_ ≈1, where the benefit of additional fibers in preventing detachment is largest. Beyond this point, binding becomes too weak for multivalency to be effective, and *N*^∗^ drops abruptly to 1, as repeated landing attempts become more efficient than surface exploration.

Geometric parameters enter through the same balance. Increasing fiber reach *r*_*s*_ increases *D*_eff_ and reduces ⟨*T*_hit_ (*M*) ⟩, shifting the optimal *N* to larger values. Increasing target size *r*_*t*_ increases both the probability of direct landing *p*_*l*_ and the surface hit rate, making the single-fiber strategy more competitive and reducing the need for high valency.

The structure underlying this optimization reflects two distinct conditions; (i) whether multivalency is globally favorable at all, determined by comparing ⟨*T*_ads_ (1) ⟩ with ⟨*T*_ads_ (*N*) ⟩, and (ii) if multivalency is beneficial, how many tail fibers to use. This number is determined by the local balance between the mobility cost of adding another tail fiber and the reduced detachment and the new host burden. We analyze these global and local transition criteria separately in the Supplemental Section S7. In high density, the global comparison ⟨*T*_ads_ (*N*) ⟩ with ⟨*T*_ads_ (1) ⟩, remains unfavorable over most of the parameter space, and the efficiency of repeated landing is not overcome by adding tail fibers. But in a sparse environment, both local and global criteria favor an increase in *N*, leading to an intermediate optimal *N* value, as shown in Fig S4.

### Effect of thermal fluctuation on adsorption rate

In addition to attachment–detachment dynamics, thermal fluctuations of the phage center-of-mass (COM) introduce an additional mechanism for local exploration of the bacterial surface (Figure 2 inset; See Table 2 for definitions.). Even when the set of attached tail fibers remains unchanged, the COM undergoes stochastic motion due to thermal forces, leading to a finite exploration region around its mean position. When we assume that each tail fiber is a flexible polymer of length *l* with the Kuhn length *κ*, approximated as a Gaussian chain with *M* = *l*/*κ* segments, the estimated effective exploration radius *r*_exp_ with *n* tail fibers out of *N* attached on the cell surface (1 ≤ *n* ≤ *N*) is given by (derivation in Supplemental Section S2)

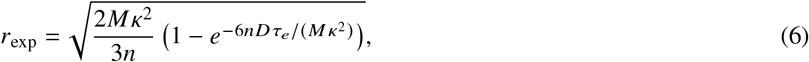

where *D* is the 3D diffusion constant of the free phage and τ_*e*_ = 1/ [(*N* −*n*) *k*_on_ +*nk*_off_] is the average time until the number of the attached tail fibers changes.

**Table 2:** Definitions and roles of the radii used in the model.

| Symbol | Definition | Interpretation |
| --- | --- | --- |
| $r_s$ | Tail-fiber reattachment reach | Maximum characteristic distance over which a detached tail fiber can reattach. It is related to length of tail fiber as shown in Figure 1(a). |
| $r_t$ | Physical target radius | Radius of the receptor or absorbing target in the absence of additional exploration due to thermal fluctuations. |
| $r_{\text{exp}}$ | Thermal exploration radius | Typical distance the phage center of mass fluctuates around its mean position while the same tail fibers stay attached (Equation (6)). The center of mass thus sweeps a region of radius $r_{\text{exp}}$; if this region overlaps the receptor, the phage can adsorb even when its mean position lies outside $r_t$. |
| $r_{\text{eff}}$ | Effective target radius | Effective target radius used in the model, $r_{\text{eff}} = \max(r_t, r_{\text{exp}})$ (Eq. 7): when $r_{\text{exp}} > r_t$, thermal fluctuations enlarge the target; otherwise the receptor size sets the target radius. |

This exploration radius can be interpreted as an effective enlargement of the target size, because if the target is with in this exploration region it will be detected. When *r*_exp_ < *r*_*t*_, thermal fluctuations of the COM are smaller than the receptor itself and do not enlarge the target, so the effective target radius reduces to the receptor radius, *r*_eff_ = *r*_*t*_. This can be expressed as an effective target size

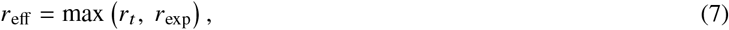

indicating that thermal fluctuations dominate the search process when *r*_exp_ > *r*_*t*_, while for smaller fluctuations, the size of the receptor sets the relevant length scale. As shown in Supplemental Figure S1, *r*_exp_ can reach tens of nanometers for realistic parameters, significantly exceeding the receptor size.

Thermal fluctuations increase the effective target size, thereby reducing the mean hitting time according to Equation (2). The magnitude of this effect depends on the mechanical properties of the tail fibers, which determine the extent of center-of-mass fluctuations and the spatial range of reattachment. Longer fibers promote a wider surface coverage, but decrease the spatial resolution of local sampling as shown in dependence of relative sites explored on *r*_*s*_ (Supplemental Figure S3).

The combined effect of the above mechanisms is shown in Figure 6, where we compare the mean adsorption time obtained using the physical target radius *r*_*t*_ and the fluctuation-enhanced effective target radius *r*_eff_ for different values of the reach of the tail-fiber *r*_*s*_. In contrast to the previous results (Figure 4(c)), we consider a larger surface (*L* = 20). For the largest tail-fiber reach (*r*_*s*_ = 50*nm*), thermal fluctuations substantially increase the effective target radius. The larger surface is therefore used to maintain a small target area fraction, consistent with the dilute-target regime assumed in the previous analysis. The physical receptor radius *r*_*t*_ is kept fixed, while *r*_eff_ is calculated from Equation (7) for each value of *r*_*s*_. The number of polymer segments is also kept fixed (*M* = 2), and since *M* = 2, increasing *r*_*s*_ corresponds to longer, but mechanically stiffer, tail fibers.

**Figure 6:**
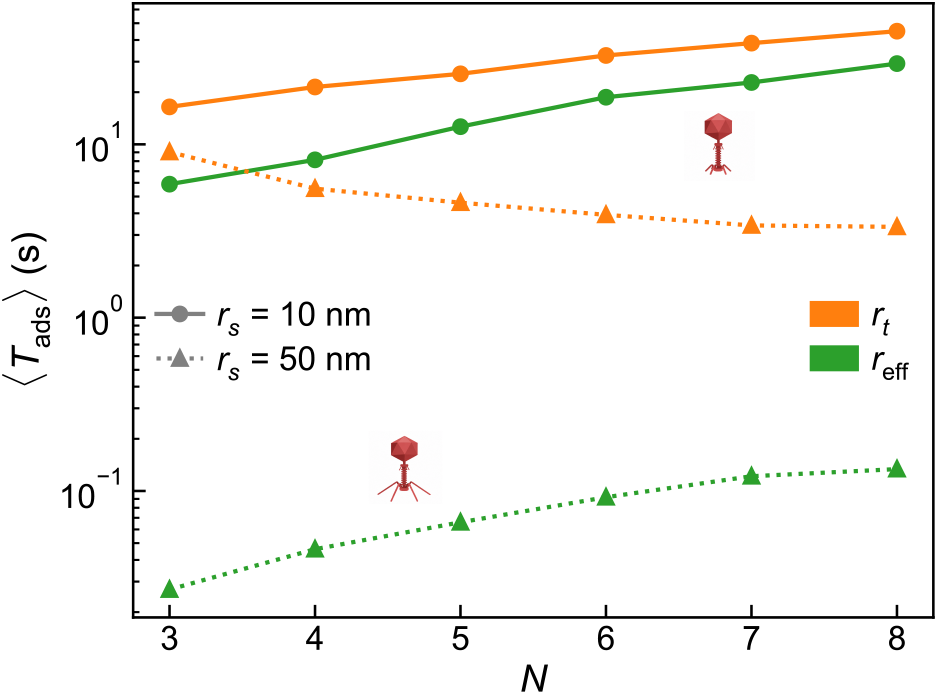
Effect of thermal fluctuations on phage adsorption: Mean adsorption time ⟨*T*_ads_⟩ as a function of tail fiber number *N* for different tail fiber reach *r*_*s*_ (with fixed number of segments per tail fiber *M* = 2), comparing fixed receptor size *r*_*t*_ and an effective target size that incorporates thermal fluctuations of the phage center-of-mass (*r*_eff_). Thermal motion increases the effective search radius and enhances target encounter probability, leading to reduced adsorption times.

Increasing the tail-fiber reach enhances adsorption through two complementary mechanisms. Longer fibers increase the displacement associated with each reattachment event, allowing the phage to explore a larger surface area by walking, while simultaneously increasing the amplitude of thermal fluctuations and hence the effective target radius. Consequently, incorporating thermal fluctuations reduces the adsorption time more strongly for longer tail fibers. This effect is most pronounced for *r*_*s*_ = 50*nm*, where the effective target size enhanced by fluctuation produces the largest reduction in the search time limited by diffusion. However, at higher values of *N*, the benefit of thermal fluctuations decreases because the attached tail fibers increasingly confine the center-of-mass motion, suppressing the fluctuation amplitude. These results indicate that thermal fluctuations provide a secondary, yet quantitatively relevant, contribution to adsorption efficiency by modifying the effective search geometry. The adsorption-time distributions for these cases are shown as violin plots in Supplemental Figure S2.

Overall, these results show that the physical properties of the tail fibers influence adsorption through multiple coupled mechanisms. Beyond determining the balance between surface mobility and attachment stability, the tail-fiber architecture also controls the extent of thermal exploration around the phage’s mean position. Although this contribution is secondary to the attachment–detachment dynamics in our model, it provides an additional means by which tail-fiber length and stiffness can modulate receptor search efficiency.

## CONCLUDING REMARKS

In this work, we investigate how the multivalent tail-surface interaction shapes the efficiency of bacteriophage adsorption by explicitly modeling the surface search as a stochastic walk. Rather than treating adsorption as a single kinetic step, we considered the physical process by which a phage explores the bacterial surface before committing to infection. This description is consistent with experimental observations that phages such as T4 undergo a reversible stepwise motion across the bacterial surface prior to irreversible adsorption (10, 11, 22). By combining stochastic simulations with analytical theory, we show that adsorption efficiency depends on the interplay between attachment, thermal vibrations, random walk by tail fibers, and stochastic resetting after complete detachment. Our results thereby link microscopic tail-fiber dynamics to the efficiency of the adsorption process.

A central outcome is that increasing the number of tail fibers does not monotonically improve adsorption. Although additional attachment points stabilize the phage location on the surface and reduce early detachment, they also restrict motion. This trade-off leads to an optimal valency, at which exploration is fast enough to locate receptors, while attachment remains sufficiently stable to avoid frequent detachment from the surface. Beyond this point, further increasing the tail fibers becomes counterproductive, as mobility is strongly suppressed and surface search slows nonlinearly. The above effects can be compensated by varying kinetic parameters *k*_on_ and *k*_off_ (Figure 5) or the tail fiber length *r*_*s*_ (Supplemental Figure S3 and Figure 6).

The dependence of the optimal adsorption strategy on environmental conditions further emphasizes that the balance between attachment stability and surface exploration is context dependent. When bacterial encounters are rare, prolonged attachment becomes advantageous because complete detachment is followed by another three-dimensional host search, a process that has long been recognized as limiting phage adsorption under low host abundance (9). In this regime a higher number of tail fibers improves performance by preventing loss of contact. In contrast, in dense environments when encounters are frequent, excessive attachment becomes a liability, and faster exploration dominates. Although the present model focuses on the microscopic surface-search process, it suggests that tail fiber number, geometry, and attachment–detachment kinetics collectively determine the efficiency of adsorption and therefore may contribute to variation in ecologically important adsorption rate (4, 6, 62).

The balance of attachment and exploration also speaks to adsorption specificity. Adsorption and productive infection are not equivalent processes, as receptor recognition must be followed by irreversible attachment and DNA delivery (15, 31). In multi-species environments, if the tail fiber associates with a non-host cell surface, the time spent on searching on those cells will be a cost for the phage, particularly for highly multivalent phages with longer surface exploration times. For example, *Clostridium difficile* phages demonstrate reversible adsorption to bacterial cells, where irreversible binding remains low, either due to the absence of correct receptors or the failure to proceed to irreversible binding and infection fails (16). Since detachment time increases with increasing tail fiber number, this represents an additional evolutionary cost of multivalency, distinct from the mobility–stability trade-off governing adsorption on the correct host.

Overall, our model provides a possible physical basis for understanding why diverse tail architectures can coexist (Table 3) (31). Rather than identifying a single optimum, the parameter space explored in Figure 5 and Figure 6 reveals that different combinations of tail-fiber number, fiber reach, and binding kinetics can achieve comparable adsorption efficiency. Such diversity may reflect adaptation to different ecological environments and infection strategies, where variations in host interactions and environmental sensing can favor distinct tail architectures (63, 64). Consequently, phages with structurally distinct tails may achieve similar adsorption performance through different combinations of physical parameters.

**Table 3:** Representative examples of phages from different ecological environments illustrating how tail-fiber multiplicity aligns qualitatively with the mobility–stability trade-offs predicted by our model. These examples are not exhaustive but demonstrate the diversity of architectures observed in nature.

| Phage / family | Host bacteria | Environment (typical) | Number of tail fibers | Fiber length (nm) | Refs |
| --- | --- | --- | --- | --- | --- |
| $\lambda$ PaPa (Siphoviridae) | *E. coli* | Laboratory | 1 (central tail fiber) | $\approx 30$ | (49–51) |
| T4 (Myoviridae) | *E. coli* | Gut / Waste | 6 long + 6 short | $\approx 144$ (long) | (27, 52) |
| T7 (Podoviridae) | *E. coli* | Gut / Waste | 6 | $\approx 32$ | (26, 53, 54) |
| SPP1 (Siphoviridae) | *B. subtilis* | Soil | 1 | $\approx 31$ | (23, 55) |
| T5 (Demereciviridae) | *E. coli* | Sewage / Freshwater | 3 (L-shaped) | $\approx 77$ | (24, 56) |
| S-PM2 (Myoviridae) | *Synechococcus* | Ocean | 6 | unknown | (57) |
| CBA120 (Kuttervirus) | *E. coli* O157:H7 | Food / Clinical | 4 (tailspikes, branched) | $\approx 16$ (TSP2) | (58, 59) |
| ΦcrAss001 (Crassvirales) | *Bacteroides intestinalis* | Human gut | 12 | unknown | (60) |
| ϕKp24 (jumbo podovirus) | *K. pneumoniae* | Clinical / Sewage | 14 (branched) | 150–170 (whole complex) | (61) |

Thermal fluctuations of phage tail fiber provide one such mechanism; the flexibility of tail fibers allows Brownian motion to sample a larger region around a given attachment configuration, effectively increasing the accessible target size during each binding state (65, 66). Similar entropic effects are well known in multivalent ligand receptor systems, where thermal fluctuations increase configurational sampling while additional binding interactions progressively restrict molecular motion (17, 67). In the present model these competing effects are reflected in the dependence of effective target size on both tail fiber length and valency Figure 6, providing a physical mechanism by which thermal fluctuation at tail fiber scale affect adsorption efficiency independently of discrete attachment detachment steps.

More broadly, this places phage adsorption within the class of stochastic search problems governed by intermittent motion and resetting (35, 36, 68, 69), where efficiency emerges from balancing surface-bound exploration against bulk excursions rather than from optimizing any single parameter. The multivalent geometry of tail fibers instantiates this balance in a way that is formally analogous to superselectivity in synthetic multivalent systems (17, 67) and to the processivity trade-offs of biological walkers (70, 71), these are the contexts in which the tension between attachment stability and mobility is a recurring design constraint.

Several simplifying assumptions should be noted. The bacterial surface was treated as homogeneous, with static, non-overlapping receptors; in reality, receptor clustering, surface heterogeneity, and cell shape are likely to further modulate search dynamics (32, 33). We also neglected possible feedback between partial attachment and later infection steps, such as conformational changes that stabilize binding or trigger irreversible adsorption. Such mechanisms are known to occur in some phages (15), and while incorporating them would likely shift quantitative predictions, we expect the qualitative trade-off between stability and mobility to persist, since it follows from the geometry of multivalent attachment rather than from any specific molecular detail.

In summary, this work shows that bacteriophage adsorption efficiency is not determined solely by molecular affinity or fiber number, but by a balance between attachment stability and spatial exploration. The existence of an optimal degree of multivalency offers a possible physical explanation for the diversity of phage tail architectures, highlighting how geometric constraints influence viral infection strategies.

## DATA AVAILABILITY

The simulation code and the data supporting the findings of this study are publicly available at https://github.com/anjali-yaadv/Phage-tail-fiber-optimization.

## DECLARATION OF GENERATIVE AI AND AI-ASSISTED TECHNOLOGIES IN THE MANUSCRIPT PREPARATION PROCESS

During the preparation of this work, the author(s) used ChatGPT (OpenAI) for paraphrasing selected sentences to improve readability and for literature searches to identify relevant publications. The author(s) reviewed and edited the output as needed and take full responsibility for the content of the published article.

## AUTHOR CONTRIBUTION

A.Y., K.S., and N.M. designed the research, developed the stochastic model and the analytical theory. A.Y. performed the simulations. A.Y. analyzed the simulation data. All authors contributed to writing the paper.

## ACKNOWLEDGMENTS

This research was funded by the Novo Nordisk Foundation (NNF21OC0068775) and the Danish National Research Foundation (grant no. DNRF170).

