## Supplementary text for "Multivalent Surface Search Dynamics Shape Bacteriophage Adsorption Efficiency: A Stochastic Model of Tail Fiber Optimization"

### SUPPLEMENTAL MATERIAL

### S1 RATE EQUATION FORM OF RENEWAL EQUATION

The renewal equation (Equation (1)) holds for arbitrary distributions $P_h(t)$, $P_d(t)$ and $P_s(t)$. Under the exponential approximation used for the analytical results (Supplemental Section S6), it is equivalent to a set of linear rate equations. While the phage remains on the surface, adsorption and detachment are competing Poisson processes with rates $\lambda_h$ and $\lambda_d$, with defective densities $P_h(t) = \lambda_h e^{-(\lambda_h+\lambda_d)t}$ and $P_d(t) = \lambda_d e^{-(\lambda_h+\lambda_d)t}$ (Supplemental Section S6), and the bulk search is exponential, $P_s(t) = \eta_B e^{-\eta_B t}$, with mean $1/\eta_B$.

Let $S(t)$ be the probability that the phage is on the cell surface and searching for a receptor, $B(t)$ the probability that it is in the bulk after complete detachment, and $A(t)$ the probability that it has adsorbed by time $t$. As in the main text, at $t = 0$ the phage starts on the cell surface with one of the tail fibers attached, $S(0) = 1$, $B(0) = A(0) = 0$. Then

$$\frac{dS}{dt} = -\left[\lambda_h + (1-p_{\rm r})\lambda_d + p_{\rm r} p_{\rm l} \lambda_d\right] S + (1-p_{\rm l})\,\eta_B\, B, \tag{S1}$$

$$\frac{dB}{dt} = (1-p_{\rm r})\,\lambda_d\, S - \eta_B\, B, \tag{S2}$$

$$\frac{dA}{dt} = (\lambda_h + p_{\rm r} p_{\rm l} \lambda_d)\, S + p_{\rm l}\,\eta_B\, B. \tag{S3}$$

Each term corresponds to a term of Equation (1). A phage on the surface finds a receptor with rate $\lambda_h$ or undergoes complete detachment with rate $\lambda_d$. After detachment, with probability $p_{\rm r}$ it returns to the same cell without a bulk-search delay; with probability $1 - p_{\rm r}$ it enters the bulk, from which it lands on a cell with rate $\eta_B$. In either case, the landing leads to immediate adsorption with probability $p_{\rm l}$ (the $\delta$-term in $K(t)$), and with probability $1 - p_{\rm l}$ the adsorption process restarts on the surface (the $P_{\rm ads}$ term in $K(t)$). Total probability is conserved, $S + B + A = 1$.

The adsorption-time distribution is the rate at which probability enters the adsorbed state,

$$P_{\rm ads}(t) = \frac{dA}{dt}, \tag{S4}$$

and coincides with the solution of Equation (1). The equivalence follows because a convolution with an exponential kernel, $y(t) = \int_0^t \lambda e^{-\lambda(t-\tau)} x(\tau)\, d\tau$, satisfies $dy/dt = \lambda x - \lambda y$, so each convolution in Equation (1) is replaced by a linear rate term.

Integrating Equations (S1) and (S2) from 0 to $\infty$ gives the mean times spent on the surface, $\tau_{\rm surf} = 1/(\lambda_h + p_{\rm l}\lambda_d)$, and in the bulk, $\tau_{\rm bulk} = (1 - p_{\rm r})\lambda_d\, \tau_{\rm surf}/\eta_B$, so that

$$\langle T(N)\rangle = \tau_{\rm surf} + \tau_{\rm bulk} = \frac{1 + \lambda_d(N)\,(1-p_{\rm r})/\eta_B}{\lambda_h(N) + p_{\rm l}\lambda_d(N)}, \tag{S5}$$

identical to the result of Supplemental Section S6 and to Equation (4). For non-exponential waiting times, such as the full first-passage-time distribution of the birth–death detachment process (Supplemental Section S4), no such finite set of rate equations exists, and the renewal form (Equation (1)) is required.

### S2 THE EFFECTIVE SEARCH RADIUS ESTIMATE

When the tail fibers are attached to the surface of a bacterium, the center of mass is expected to explore some distance due to thermal fluctuations. We estimate the order of magnitude of this range by assuming the following:

(i) A tail fiber is a flexible polymer of length $\ell$ and the Kuhn length $\kappa$, and we approximate it as a gaussian chain of bond length $\kappa$ with $M = \ell/\kappa$ segments.

(ii) $n$ tail fibers out of $N$ are attached on a cell surface. We consider the situation where $1 \le n \le N$.

(iii) When the center of mass coincides with the center of the $n$ attached tail fibers, each tail fiber is at its natural (unstretched/uncontracted) length, and the attachment points are distributed symmetrically on a circle around the center of mass.

With the Gaussian chain assumption (i), the distribution of three dimensional end-to-end distance $R$ of the tail fibers obeys

$$\Phi(R) = \left(\frac{3}{2\pi M\kappa^2}\right)^{3/2} \exp\left(-\frac{3R^2}{2M\kappa^2}\right).$$

This gives the free energy

$$F(R) = -k_B T \ln \Phi(R) = \frac{3k_B T}{2M\kappa^2} R^2 + \text{constant}.$$

with $k_B$ and $T$ being the Boltzmann constant and the temperature, respectively. Equating the $R$-dependent part of the energy to that of a spring, we obtain the effective spring constant to be

$$k = \frac{3k_B T}{M\kappa^2}.$$

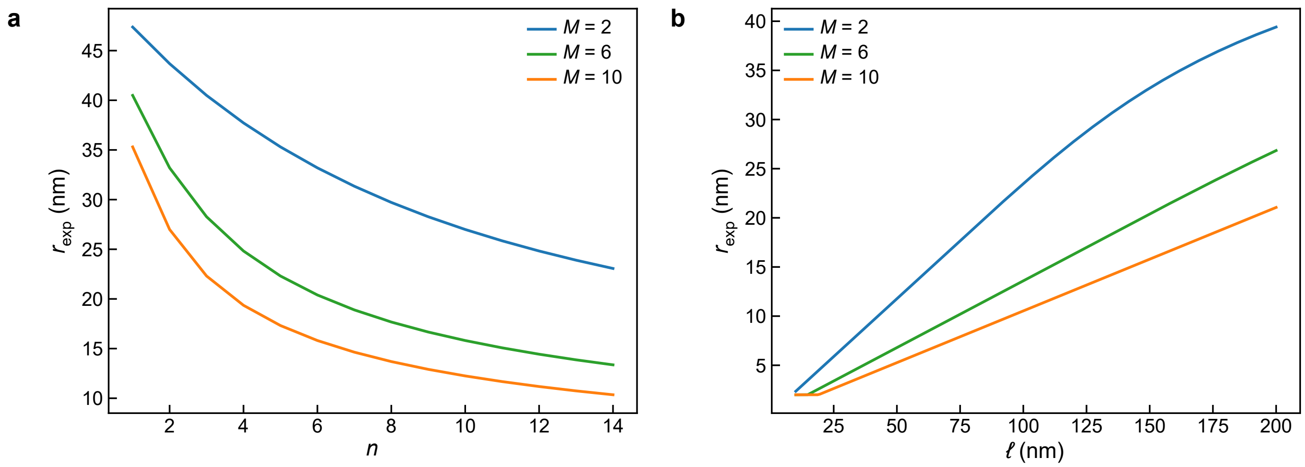


Figure S1: The effective search radius $r_{\text{exp}}$ represents the expected center-of-mass (COM) position of the phage under thermal fluctuations while attached to the surface. Even when tail fiber attachments remain fixed, thermal motion induces fluctuations in the COM, effectively broadening the region over which a receptor can be detected. As a result, $r_{\text{exp}}$ can exceed the intrinsic receptor radius $r_t$, leading to an increased effective target size. (a) $r_{\text{exp}}$ as a function of the number of attached tail fibers $n$ for different fiber softness (parameterized by $M$). (b) $r_{\text{exp}}$ as a function of tail fiber length for different values of $M$. Conversely, for small fiber reach $r_s$, fluctuations are suppressed and $r_{\text{exp}} \approx r_t$, so the effective search area is limited by the physical receptor size. Default parameters: $D = 4\,\mu\text{m}^2/\text{s}$, $\tau_e = (10^{-3}/N)s$
, $N = 6$, $n = 6$, and $l = 150\,\text{nm}$.

Next, we evaluate the fluctuation of the center of mass under the assumption (iii). The dislocation of center of mass from the equilibrium position, $\mathbf{X}_c$, will obey the following Langevin equation

$$\frac{d\mathbf{X}_c}{dt} = -\mu n k \mathbf{X}_c + \xi,$$

where the random force $\xi$ is a Gaussian white noise, amplitude determined by the diffusion constant of the center of mass $D$, and $\mu$ is the mobility of the center of mass. The Einstein's relation ensures

$$D = \mu k_B T.$$

Assuming $\mathbf{X}_c(0) = 0$ at time $t = 0$, the variance of the position of the center of mass projected to the surface plain at time $t$ is given by

$$\langle \Delta r(t)^2 \rangle = \langle X_c^2 + Y_c^2 \rangle = \frac{2D}{\mu n k}\left(1 - e^{-2nk\mu t}\right) = \frac{2M\kappa^2}{3n}\left(1 - e^{-6nDt/(M\kappa^2)}\right),$$

where in the last equality expression of the spring constant and Einstein's relation were used.

This setup is valid until one of the tail fiber detaches or newly attaches. Average time $\tau_e$ until one of these events to happen is given by

$$\tau_e = \left[(N-n)k_{\text{on}} + nk_{\text{off}}\right]^{-1}.$$

The typical radius $r_{exp}$ that the center of mass explore in this duration is given by

$$r_{exp} = \sqrt{\langle \Delta r(\tau_e)^2 \rangle}.$$

Now, we will estimate the plausible range of $r_{exp}$. For simplicity, we assume $k_{\text{on}} = k_{\text{off}} = 10^3/\text{s}$, giving $\tau_e = (10^{-3}/N)\text{s}$, and the phage diffusion constant is about $D = 4\mu\text{m}^2/\text{s}$ (for phage T4 in water).

For a T4 phage, the $N = 6$ long tail fiber is about $\ell \sim 0.15\mu$ m is known to be relatively stiff except for the hinge region around the middle. Assuming then $M = 2$ (hence $\kappa = 0.075\mu\text{m}$), we have

$$r_{exp} \approx \sqrt{\frac{4}{3n}\left(1 - e^{-0.35n}\right)\kappa},$$

which spans 32 nm for $n = 6$ and 43 nm for $n = 2$.

If we assume the same geometry but significantly softer fibers of $M = 6$ (hence $\kappa = 0.025\mu\text{m}$), we have

$$r_{exp} \approx \sqrt{\frac{4}{n}\left(1 - e^{-6.4n}\right)\kappa},$$

giving 20nm ($n = 6$) to 30nm ($n = 2$).

On the other hand, if we assume a short and soft tail fiber of e.g. 30nm, with $M = 6$ (hence $\kappa = 0.005\mu\text{m}$), $r_{exp} \approx 4 \sim 7\text{nm}$.

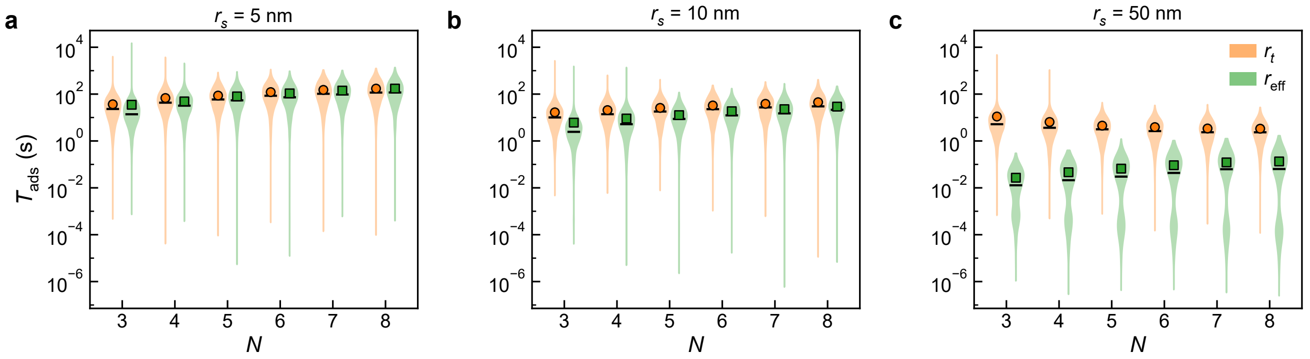


Figure S2: (a) $r_s = 5$ nm. (b) $r_s = 10$ nm (c) $r_s = 50$ nm. Distribution of total adsorption times $P_{\text{ads}}(t)$ for two scenarios: fixed target radius ($r_t$) and dynamically varying effective target radius based on the number of attached fibers, given by eq S13.The black line shows meadian of adsorption time and square and circle show mean of adsoprtion time respectively for fixed and variable $r_t$

### S3 TRADEOFF DUE TO $r_s$

To isolate the role of tail-fiber reach, we varied the span radius $r_s$ while keeping all other parameters fixed at their default values, including $r_t = 0.2$. Increasing $r_s$ increases the step size associated with each attachment–detachment event and therefore enhances the effective surface mobility of the phage, consistent with the scaling $D_{\text{eff}} \propto r_s^2$, shown in Figure S3(a). Figure S3(b) shows the corresponding total adsorption-time distributions for different values of $r_s$. As $r_s$ increases, the distribution shifts systematically toward shorter times, indicating more efficient adsorption. This trend is consistent with the interpretation that longer reattachment reach allows the phage center of mass to explore the surface more rapidly, and hence reducing the time required to encounter the target. For the default receptor size considered here, $r_t = 0.2$ and $N = 6$, we do not observe a competing penalty from increased step size. Instead, the dominant effect of increasing $r_s$ is to accelerate surface exploration. Thus, within this parameter range, larger fiber reach monotonically improves adsorption efficiency.

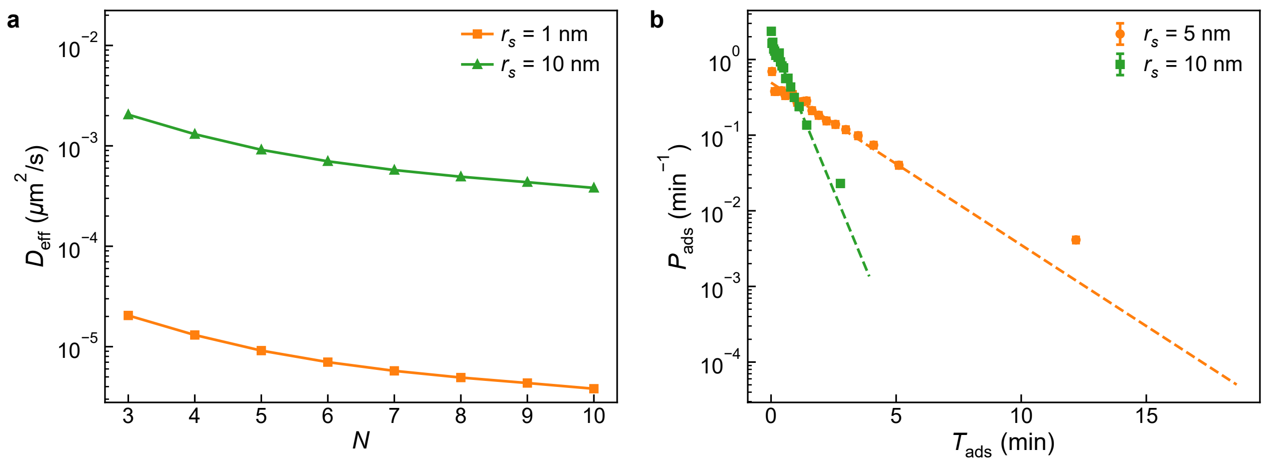


Figure S3: a) Effect of fiber reach $r_s$ on diffusion constant with default parameters, b) Effect of fiber reach $r_s$ on adsorption time.

To quantify surface exploration efficiency, the surface of the bacteria was discretized into a square lattice $m \times m$ and the number of distinct lattice sites visited by the center of mass within a fixed time window. Gillespie simulations are run in regime $k_{\text{off}} << k_{\text{on}}$, ensuring that the phage does not detach from the surface. Only trajectories where the phage remains attached to the surface throughout the time window were used for analysis. Figure 5(c) shows the number of unique surface lattice sites explored as a function of the number of tail fibers N for different tail fiber span radius, $r_s$.

### S4 DETACHMENT RATE $\lambda_d$ FROM A BIRTH–DEATH MASTER EQUATION (START AT ONE TAIL FIBER ATTACHED)

**Model** We consider an adhesion complex with up to $N$ independent bonds. Let $n \in \{0, 1, \ldots, N\}$ denote the number of attached bonds. State $n = 0$ is absorbing (fully detached). From state $n \geq 1$,

$$\lambda_n = k_{\text{off}}\, n, \qquad \mu_n = k_{\text{on}}\,(N - n)$$

are the total *detachment* ($n \to n - 1$) and *reattachment* ($n \to n + 1$) rates, respectively; at the top boundary $\mu_N = 0$. We seek the mean first-passage time (MFPT) to detachment,

$$m_n \;=\; \mathbb{E}[\text{time to hit } 0 \mid \text{start at } n],$$

and, in particular, $m_1$ (start from a single attached bond). The effective detachment rate used in the renewal equation is

$$\lambda_d \;:=\; \frac{1}{m_1}.$$

For $1 \leq n \leq N - 1$ the standard backward equations read

$$-1 \;=\; -(\lambda_n + \mu_n)\, m_n \;+\; \lambda_n m_{n-1} \;+\; \mu_n m_{n+1},$$

with boundary conditions

$$m_0 = 0, \qquad -1 \;=\; -\lambda_N m_N + \lambda_N m_{N-1} \quad (\text{since } \mu_N = 0). \tag{S6}$$

**First-difference trick.** Define the first differences $\Delta_n := m_n - m_{n-1}$ for $n \geq 1$. Subtracting adjacent equations gives the telescoping form

$$\mu_n \, \Delta_{n+1} \; - \; \lambda_n \, \Delta_n \; = \; -1 \qquad (1 \leq n \leq N-1),$$

and at the top boundary, from Equation (S6),

$$\Delta_N \; = \; m_N - m_{N-1} \; = \; \frac{1}{\lambda_N}.$$

We can now iterate *downwards*. Rewriting

$$\Delta_n \; = \; \frac{\mu_n}{\lambda_n} \, \Delta_{n+1} \; + \; \frac{1}{\lambda_n} \qquad (1 \leq n \leq N-1),$$

and unrolling from $n = N-1$ to $n = 1$ yields the closed form

$$\Delta_k \; = \; \sum_{j=k}^{N} \frac{1}{\lambda_j} \prod_{r=k}^{j-1} \frac{\mu_r}{\lambda_r}, \qquad 1 \leq k \leq N,$$

with the convention that an empty product equals 1. Finally,

$$m_1 \; = \; \sum_{k=1}^{1} \Delta_k \; = \; \Delta_1 \; = \; \sum_{j=1}^{N} \frac{1}{\lambda_j} \prod_{r=1}^{j-1} \frac{\mu_r}{\lambda_r}. \tag{S7}$$

**Evaluate for our rates.** Insert $\lambda_n = k_{\text{off}} \, n$ and $\mu_n = k_{\text{on}} \, (N-n)$ into Equation (S7). Let $a := k_{\text{off}}$, $b := k_{\text{on}}$, and $x := b/a$. Then

$$\frac{\mu_r}{\lambda_r} \; = \; \frac{b(N-r)}{a \, r} \quad \Rightarrow \quad \prod_{r=1}^{j-1} \frac{\mu_r}{\lambda_r} \; = \; \left(\frac{b}{a}\right)^{j-1} \frac{(N-1)!}{(N-j)! \, j!} \; = \; x^{j-1} \, \frac{1}{j} \binom{N-1}{j-1}.$$

Moreover, $\lambda_j = a \, j$. Therefore,

$$m_1 = \sum_{j=1}^{N} \frac{1}{a \, j} x^{j-1} \, \frac{1}{j} \binom{N-1}{j-1} \, j \; = \; \frac{1}{a} \sum_{j=1}^{N} \frac{1}{j} \binom{N-1}{j-1} x^{j-1}. \tag{S8}$$

Use the binomial identity $\frac{1}{j}\binom{N-1}{j-1} = \frac{1}{N}\binom{N}{j}$ to obtain

$$m_1 = \frac{1}{a} \cdot \frac{1}{N} \sum_{j=1}^{N} \binom{N}{j} x^{j-1} = \frac{1}{a} \cdot \frac{1}{Nx} \sum_{j=1}^{N} \binom{N}{j} x^{j} \tag{S9}$$

$$= \frac{1}{a} \cdot \frac{(1+x)^N - 1}{Nx} = \frac{(1+x)^N - 1}{N \, b} \qquad \left(x = \frac{b}{a}\right). \tag{S10}$$

**Effective detachment rate.** The detachment hazard used in the renewal equation is the inverse MFPT,

$$\boxed{\lambda_d \; = \; \frac{1}{m_1} \; = \; \frac{N \, k_{\text{on}}}{\left(1 + \frac{k_{\text{on}}}{k_{\text{off}}}\right)^N - 1}}.$$

### S5 THEORETICAL ANALYSIS OF SURFACE SEARCH HIT-TIME DISTRIBUTION

#### Mean First Passage Time on a Periodic Surface

The phage center-of-mass undergoes 2D Brownian motion with effective diffusion constant $D_{\text{eff}}$ on a periodic domain $T^2 = [0, L] \times [0, L]$ until encountering a circular absorbing target of radius $r_t \ll L$. The mean first passage time $\tau(\mathbf{r})$ from position $\mathbf{r}$ satisfies:

$$D_{\text{eff}}\nabla^2\tau(\mathbf{r}) + 1 = 0 \quad \text{in } \Omega = T^2 \setminus B_{r_t} \tag{S11}$$

with boundary conditions $\tau = 0$ on $\partial B_{r_t}$ and periodic conditions on $\partial T^2$.

The uniform source in Equation (S11) requires a neutralizing background on the periodic domain. We construct Green's function $G(\mathbf{x}, \mathbf{y})$ satisfying:

$$-D_{\text{eff}}\nabla_x^2 G(\mathbf{x}, \mathbf{y}) = \delta_{T^2}(\mathbf{x} - \mathbf{y}) - \frac{1}{L^2} \tag{S12}$$

Using Fourier basis $\phi_{mn}(\mathbf{x}) = L^{-1}e^{i\mathbf{k}_{mn}\cdot\mathbf{x}}$ with $\mathbf{k}_{mn} = \frac{2\pi}{L}(m, n)$, and excluding the $(0, 0)$ mode for neutrality:

$$G(\mathbf{x}, \mathbf{y}) = \frac{1}{4\pi^2 D_{\text{eff}}} \sum_{(m,n)\neq(0,0)} \frac{1}{m^2 + n^2} e^{i\mathbf{k}_{mn}\cdot(\mathbf{x}-\mathbf{y})} \tag{S13}$$

For small $r = |\mathbf{x} - \mathbf{y}|$, the Green's function has the local behavior:

$$G(\mathbf{x}, \mathbf{y}) = -\frac{1}{2\pi D_{\text{eff}}} \ln r + R_{\text{torus}} + O(r^2) \tag{S14}$$

We construct the outer solution $\tau_{\text{out}}(\mathbf{x}) = \bar{\tau} - L^2 G(\mathbf{x}, \mathbf{x}_0)$ and match with the inner solution $\tau_{\text{in}}(r) = A\ln(r/r_t)$ in the region $r_t \ll r \ll L$. Matching coefficients yields:

$$\bar{\tau} = \frac{L^2}{2\pi D_{\text{eff}}}\left[\ln\left(\frac{L}{r_t}\right) + C_{\text{torus}}\right] \tag{S15}$$

where $C_{\text{torus}} = 2\pi D_{\text{eff}} R_{\text{torus}} - \ln L$ is the geometric constant.

Under rapid mixing conditions, the survival probability decays exponentially with rate:

$$\lambda_h = \frac{1}{\langle T_{\text{hit}}\rangle} = \frac{2\pi D_{\text{eff}}}{L^2(\ln(L/r_t) + C_{\text{torus}})} \tag{S16}$$

leading to the exponential distribution $P_h(t) = \lambda e^{-\lambda t}$ observed in simulations (Figure 3(a).

Using the scaling obtained for $D_{eff}$ from the diffusion analysis in the main text, we get mean hit time,

$$\langle T_h\rangle = \frac{L^2}{2\pi k_{\text{off}} r_s^2}\left(\ln\left(\frac{L}{r_t}\right) + C_{\text{torus}}\right) N^{\alpha}.$$

Collecting the geometry- and parameter-dependent factors into a constant prefactor,

$$A \equiv \frac{L^2}{2\pi k_{\text{off}} r_s^2} \ln\left(\frac{L}{r_t}\right),$$

For $n_t$ well-separated targets with area fraction $\phi = n_t \pi r_t^2 / L^2 \ll 1$:

$$\langle T_{\text{hit}}\rangle \approx \frac{L^2}{4\pi D_{\text{eff}} n_t r_t^2}\left[\ln\left(\frac{L^2}{n_t \pi r_t^2}\right) + \gamma - \frac{1}{4}\right] \tag{S17}$$

with $\gamma = 0.5772$.

#### Validation and Parameter Fitting

The theoretical rate $\lambda$ (Equation (S16)) requires the effective diffusion constant $D_{\text{eff}}$ measured from MSD analysis (Fig. 1C) and the geometric constant $C_{\text{torus}}$. Fitting simulation data to theory yields $C_{\text{torus}} \approx -1.2$ for our system geometry. The exponential character of $P_h(t)$ validates the rapid mixing assumption across all parameter regimes tested.

### S6 DERIVATION OF THE MEAN ADSORPTION TIME

We now derive the mean adsorption time under the same exponential approximation used in the main text. We assume that while the phage remains on the surface, adsorption and detachment are competing Poisson processes with rates $\lambda_h(N)$ and $\lambda_d(N)$, respectively. The corresponding defective densities are

$$P_h(t) = \lambda_h\, e^{-(\lambda_h+\lambda_d)t}, \qquad P_d(t) = \lambda_d\, e^{-(\lambda_h+\lambda_d)t},$$

so that

$$q_h = \int_0^\infty P_h(t)\, dt = \frac{\lambda_h}{\lambda_h + \lambda_d}, \qquad q_d = \int_0^\infty P_d(t)\, dt = \frac{\lambda_d}{\lambda_h + \lambda_d},$$

with $q_h + q_d = 1$. The mean time to the first event on the surface is therefore

$$\langle t_{\text{first}} \rangle = \frac{1}{\lambda_h + \lambda_d}.$$

Let $\langle T(N) \rangle$ denote the mean total adsorption time starting from a landing on the cell surface. Conditioning on the first event gives

$$\langle T(N) \rangle = \frac{1}{\lambda_h + \lambda_d} + q_d \, \langle T_{\text{reset}} \rangle, \tag{S18}$$

where $\langle T_{\text{reset}} \rangle$ is the mean additional time accumulated after a detachment event.

After detachment, two cases are possible. With probability $p_{\text{r}}$, the phage returns to the same cell without a bulk-search delay. With probability $1 - p_{\text{r}}$, it enters the bulk and waits a mean time $1/\eta_B$ before landing on a cell. In either case, the subsequent landing leads to immediate adsorption with probability $p_1$, and with probability $1 - p_1$ the adsorption process restarts. Hence

$$\langle T_{\text{reset}} \rangle = p_{\text{r}} \left[ p_1 \cdot 0 + (1-p_1)\langle T(N) \rangle \right] + (1-p_{\text{r}}) \left[ \frac{1}{\eta_B} + p_1 \cdot 0 + (1-p_1)\langle T(N) \rangle \right].$$

Collecting terms,

$$\langle T_{\text{reset}} \rangle = \frac{1-p_{\text{r}}}{\eta_B} + (1-p_1)\langle T(N) \rangle.$$

Substituting the expression for $\langle T_{reset} \rangle$ into Equation (S18) gives

$$\langle T(N) \rangle = \frac{1}{\lambda_h + \lambda_d} + \frac{\lambda_d}{\lambda_h + \lambda_d} \left[ \frac{1-p_{\text{r}}}{\eta_B} + (1-p_1)\langle T(N) \rangle \right].$$

Rearranging,

$$\langle T(N) \rangle \left[ 1 - \frac{\lambda_d}{\lambda_h + \lambda_d}(1-p_1) \right] = \frac{1}{\lambda_h + \lambda_d} + \frac{\lambda_d}{\lambda_h + \lambda_d}\, \frac{1-p_{\text{r}}}{\eta_B},$$

and therefore

$$\boxed{\langle T(N) \rangle = \frac{1 + \lambda_d(N)\,(1-p_{\text{r}})/\eta_B}{\lambda_h(N) + p_1 \lambda_d(N)}.}$$

Using the empirical scaling

$$\lambda_h(N) = \frac{1}{AN},$$

together with the birth–death result

$$\lambda_d(N) = \frac{N k_{\text{on}}}{(1 + k_{\text{on}}/k_{\text{off}})^N - 1},$$

we obtain

$$\boxed{\langle T(N) \rangle = \frac{AN \left[ 1 + (1-p_{\text{r}}) \dfrac{N k_{\text{on}}/\eta_B}{(1 + k_{\text{on}}/k_{\text{off}})^N - 1} \right]}{1 + AN p_1 \dfrac{N k_{\text{on}}}{(1 + k_{\text{on}}/k_{\text{off}})^N - 1}}.}$$

The effect of re-hitting is to reduce the detachment penalty by lowering the mean bulk-search delay after detachment, while the effect of direct landing is to increase the effective success probability upon each return to the surface.

The case $N = 1$ must be treated separately. In this limit, there is no multivalent walking state, and the surface-search description based on $\lambda_h(N)$ is no longer applicable. Instead, adsorption proceeds through repeated landing attempts on the cell surface.

Upon landing on a cell, adsorption occurs immediately with probability $p_{\mathrm{l}}$. If adsorption does not occur, the phage remains attached by its single tail fiber until detachment, which occurs with rate

$$\lambda_d(1) = k_{\mathrm{off}}.$$

After detachment, the phage returns to the same cell with probability $p_{\mathrm{r}}$. Otherwise, with probability $1 - p_{\mathrm{r}}$, it undergoes a bulk-search phase with mean duration $1/\eta_B$ before landing again.

Let $\langle T \rangle_{N=1}$ denote the mean adsorption time for $N = 1$, measured from a landing on the cell surface. Conditioning on the outcome of the first landing gives,

$$\langle T \rangle_{N=1} = \frac{1}{p_l}\left[\frac{1}{k_{\mathrm{off}}} + \frac{1 - p_r}{\eta_B}\right]$$

### S7 GLOBAL AND LOCAL OPTIMALS

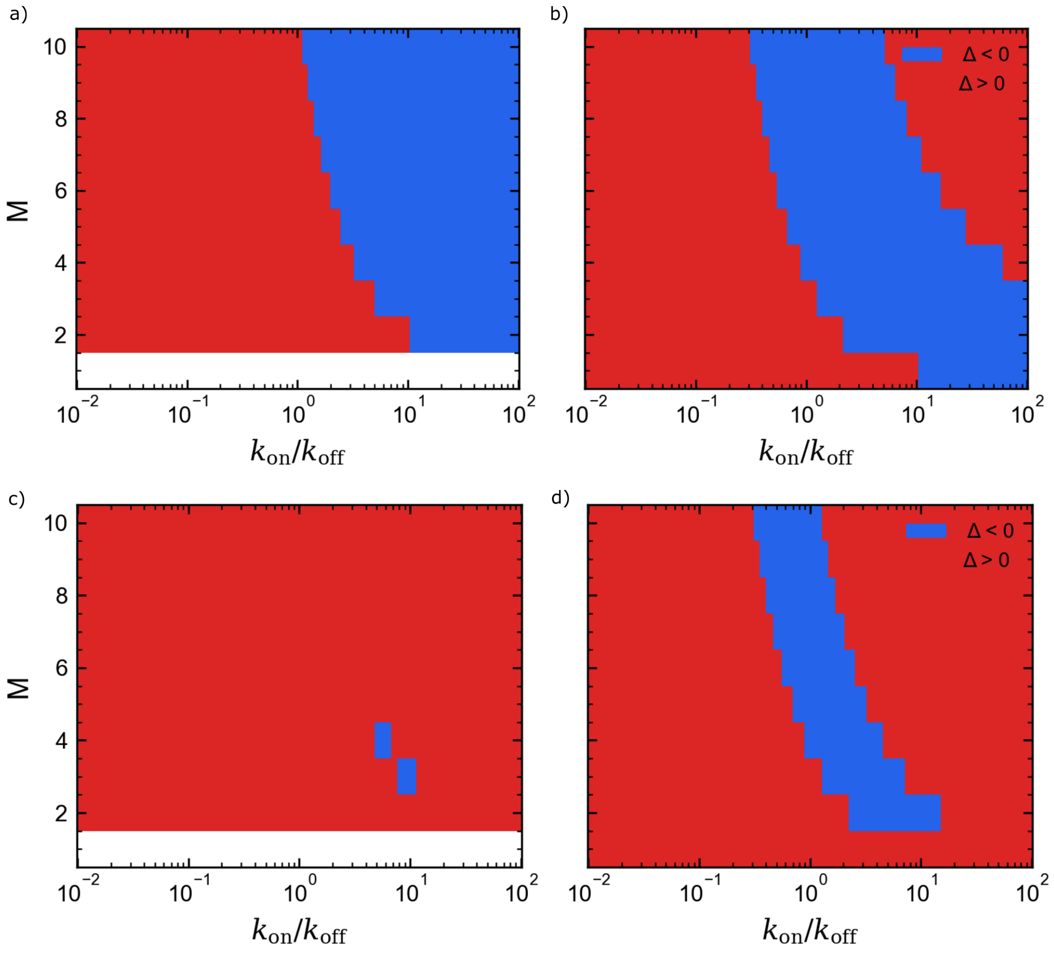


Figure S4: Sign maps of (left) the global comparison and (right) the local incremental change, shown as functions of $k_{\mathrm{on}}/k_{\mathrm{off}}$ and $M$, for (top) low-density and (bottom) high-density environments. Blue regions correspond to $\Delta < 0$ (favorable), while red regions correspond to $\Delta > 0$ (unfavorable).

To further elucidate the structure of the optimal valency landscape, we separately examine global and local transition criteria across parameter space Figure S4

The left panels show the sign of the global comparison

$$\Delta_M^{(1)} = \langle T_{\text{ads}}(M)\rangle - \langle T_{\text{ads}}(1)\rangle,$$

which determines whether an $M$-tail fiber strategy is favorable relative to the single-fiber state. The right panels show the sign of the local incremental change

$$\Delta_M^{\text{loc}} = \langle T_{\text{ads}}(M+1)\rangle - \langle T_{\text{ads}}(M)\rangle,$$

which determines whether adding one additional tail fiber is beneficial within the multivalent regime. Blue regions correspond to $\Delta < 0$ (favorable), while red regions correspond to $\Delta > 0$ (unfavorable).

It is useful to distinguish two different transition criteria. The first is single to multivalent comparison, defined by comparing the single-fiber strategy to an multivalent state $M \geq 2$:

$$\langle T_{ads}(1)\rangle = \langle T_{ads}(M)\rangle$$

This criteria determines when multivalency becomes favorable at all. For $N = 1$, adsorption reduces to repeated independent landing attempts. A successful adsorption occurs only if the phage lands directly on the receptor region with probability $p_l$, depending on receptor density, while each unsuccessful attempt costs the mean attachment time before detaching, i.e, $1/k_{\text{off}}$, plus the mean time to find another host, $(1 - pr)/\eta B$. The global condition $T_1 = T_M$ therefore compares two different search modes, repeated direct landings versus surface exploration on the cell with repeated landing. The second is local multivalent transition, comparing neighbouring multivalent state:

$$\langle T_{ads}(M)\rangle = \langle T_{ads}(M+1)\rangle, \qquad M \geq 2$$

This determines how optimum is selected once surface search with multivalency is advantageous than single tail fiber. In the limit $p_l << 1$, this can be interpreted as balance between cost and benefit of adding more tail fiber. The local transition $\langle T_{\text{ads}}(M+1)\rangle = \langle T_{\text{ads}}(M)\rangle$ can be interpreted as a balance between two competing effects:

$$\underbrace{\langle T_{\text{hit}}(M+1)\rangle - \langle T_{\text{hit}}(M)\rangle}_{\text{mobility cost}} = \underbrace{\frac{1-p_r}{\eta_B}\left[\frac{\langle T_{\text{hit}}(M)\rangle}{\langle T_{\text{det}}(M)\rangle} - \frac{\langle T_{\text{hit}}(M+1)\rangle}{\langle T_{\text{det}}(M+1)\rangle}\right]}_{\text{reduction in detachment and finding new target delay}}. \tag{S19}$$

The left-hand side represents the increase in surface search time due to reduced mobility when an additional tail fiber is added. The right-hand side captures how additional fibers reduce the delay arising from detachment events that compete with successful target finding.

The ratio $\langle T_{\text{hit}}(N)\rangle/\langle T_{\text{det}}(N)\rangle$ quantifies the relative timescales of these competing processes: when this ratio is large, detachment occurs on a faster timescale than target finding and therefore significantly contributes to the overall adsorption time. The environmental encounter rate $\eta B$ enters through the reset time $(1 - p_r)/\eta B$, and therefore sets the scale of both transition criteria. In dense environments, where $\eta B$ is large and the reset time is short, detachment is relatively inexpensive. Under these conditions, the global comparison $\langle T_{\text{ads}}(1)\rangle = \langle T_{\text{ads}}(M)\rangle$ favors small $N$, since repeated landing attempts are efficient, and the local balance is dominated by mobility costs. In contrast, in sparse environments where $\eta B$ is small and the reset time is large, detachment is costly, favoring multivalency and shifting both global and local transitions toward larger optimal $N$. In the low-density regime (high panels), the global comparison strongly favors multivalency, and the local transitions exhibit a systematic progression toward larger $M$. This reflects the increasing importance of suppressing costly detachment-reset events when the encounter rate $\eta B$ is small. In contrast, In the high-density regime (bottom panels), the global comparison remains unfavorable over most of parameter space, indicating that repeated landing attempts are already efficient and multivalency provides little advantage. Although local transitions can occasionally favor increasing $N$, these incremental improvements do not overcome the global disadvantage relative to the single-fiber strategy.

These results demonstrate that the onset of multivalency and the selection of the optimal valency are governed by distinct criteria. In particular, intermediate increases in tail fiber number can be locally unfavorable even when larger multivalent states yield a lower overall adsorption time, highlighting that global optimality is not necessarily achieved through monotonic local transitions.
